# Phylogeny of the ingoid clade (Caesalpinioideae, Fabaceae), based on nuclear and plastid data

**DOI:** 10.1101/2021.11.23.469677

**Authors:** Julia Ferm, Bertil Ståhl, Niklas Wikström, Catarina Rydin

## Abstract

We investigated generic relationships in the ingoid clade (Fabaceae) (sensu Koenen & al. 2020a), with main focus on genera with a taxonomic history in *Calliandra* s.l. of the tribe Ingeae (i.e. *Afrocalliandra, Calliandra* s.s., *Sanjappa, Thailentadopsis, Viguieranthus, Zapoteca*), and three genera of the tribe Acacieae (i.e., *Acacia, Acaciella, Senegalia*). The nuclear ribosomal ETS and ITS, and the plastid *matK, trnL-trnF* and *ycf1* DNA-regions were analysed for 246 representatives from 36 genera using maximum likelihood as implemented in IQ-tree. The results show an Ingeae–*Acacia* clade within the ingoid clade, resolved in three major clades. Clade 1 (*Calliandra* s.s. and *Afrocalliandra*) is sister to clades 2 and 3. Clade 2 comprises *Faidherbia, Sanjappa, Thailentadopsis, Viguieranthus* and *Zapoteca*. Clade 3 comprises the remaining genera of the Ingeae, plus *Acacia*. The ingoid genus *Senegalia* is excluded from the Ingeae–*Acacia* clade. *Acaciella* is sister to the remaining ingoid clade when nuclear ribosomal data is included in the analyses, but included in the Ingeae–*Acacia* clade based on plastid data. *Acacia* and perhaps also *Acaciella* are thus nested within Ingeae. Species traditionally referred to *Calliandra* (*Calliandra* s.l.) are resolved in two clades, and the “*Calliandra*-pod” has apparently evolved independently several times.

## Introduction

The legume family, Fabaceae, is globally distributed and consists of approximately 19 500 species in about 750 genera (Lewis & al. 2005). Traditionally, three legume subfamilies (i.e. Mimosoideae, Papilinoideae and Caesalpinioideae) have been recognized based on morphological characters. However, several studies have shown that while both Mimosoideae and Papilinoideae are monophyletic, the Mimosoideae are nested within Caesalpinioideae, making Caesalpinioideae paraphyletic (Wojciechowski & al. 2004; Bruneau & al. 2008; LPWG 2013; LPWG 2017). Thus, the family classification has recently been revised to include six subfamilies, with a monophyletic mimosoid clade (the former subfamily Mimosoideae) included in Caesalpinioideae (LPWG 2017). The mimosoid clade includes four tribes: Acacieae, Ingeae, Mimoseae and the monospecific Mimozygantheae (Lewis & al. 2005). However, Acacieae, Ingeae and Mimoseae have all been shown to be non-monophyletic (Luckow & al. 2003; Miller & Seigler 2012; Kyalangalilwa & al. 2013). The Ingeae tribe is paraphyletic with regards to the genus *Acacia* Mill. of tribe Acacieae (Miller & Bayer 2001; Miller & al. 2003; Brown & al. 2008; LPWG, 2017; Ferm & al. 2019). A recent study by Koenen et al (2020a) addressed major relationships in the mimosoid clade based on massive amounts of low copy nuclear genes, and found that a *“Mimosa* clade” is sister to an “ingoid clade”, which includes the tribe Ingeae as well as the genera *Acacia, Mariosousa* Seigler & Ebinger, *Senegalia* Raf. and *Acaciella* Britton & Rose of the tribe Acacieae. Species of this ingoid clade have more than 10 stamens (Fig. 1), which in most genera are fused together into a tube. Having such synandrous flowers is the one character that traditionally has been used to distinguish species of the Ingeae tribe from other mimosoids (Bentham, 1865).

**Fig. 1.**
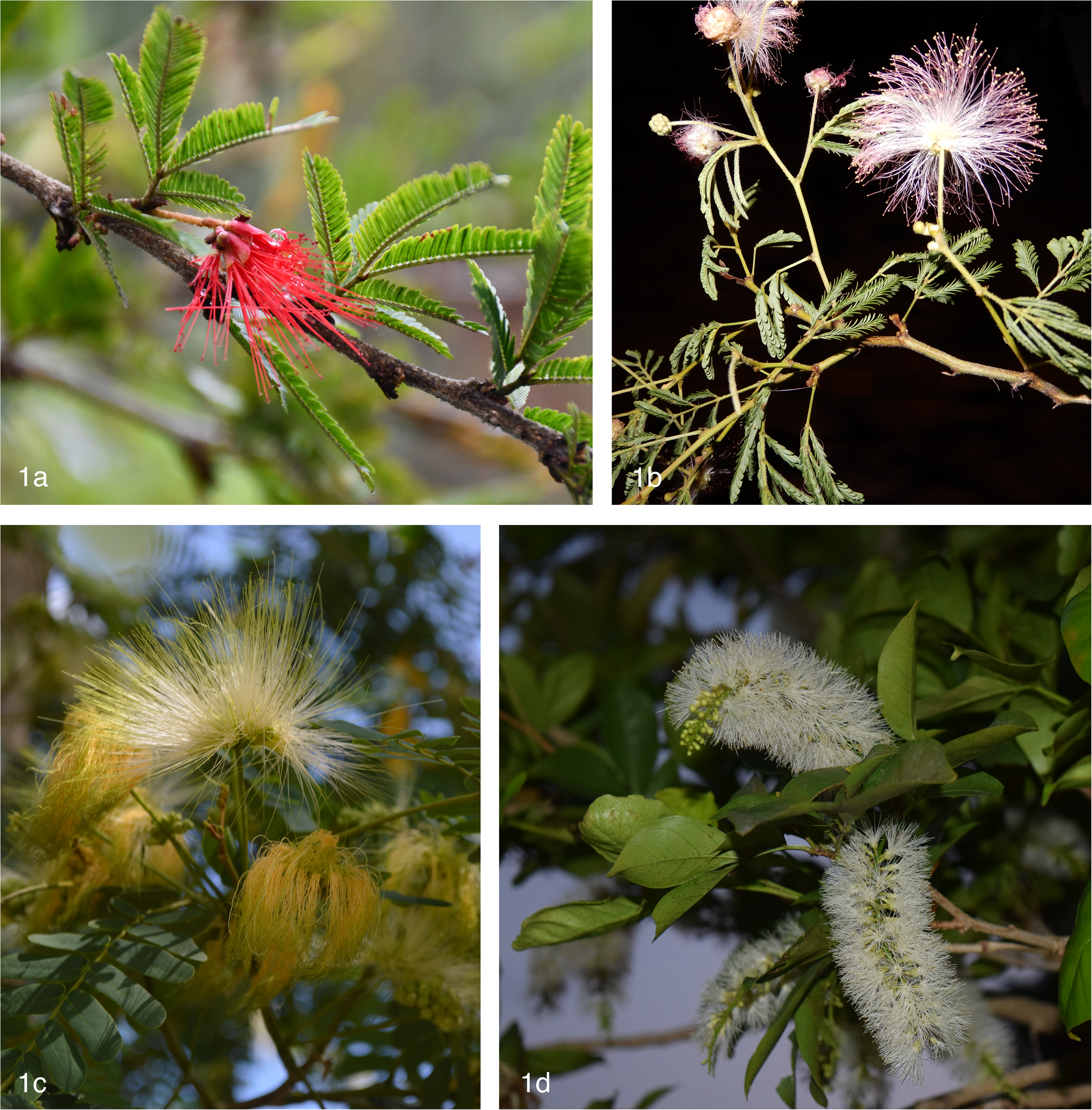
Inflorescences showing synandrous flowers of 1a. *Calliandra taxifolia* 1b. *Zapoteca caracasana* 1c. *Albizia lebbeck* 1d. *Inga laurina* Photo: Bertil Ståhl (a-b), Steve Maldonado Silvestrini (c-d).

The Ingeae tribe has a history of taxonomic instability at the generic level. The number of recognized genera has increased over the years (Bentham 1875; Nielsen 1981; for a detailed summary, see Brown 2008), with the latest formal classification including 36 genera placed in seven informal groups (Lewis & Rico Arce 2005) mainly following Barneby & Grimes (1996), who recognized 5 alliances in Ingeae. This taxonomic instability is also reflected at the species level; many species have a history of being placed in many different genera. For example, the well known rain tree *Samanea saman* (Jacq.) Merr., has been placed in nine different genera (POWO 2021). Generic circumscriptions and species taxonomy continue to change in the light of new phylogenetic discoveries, and the re-classification of Ingeae is evidently still a work in progress (e.g. Souza & al. 2013, 2016; Ferm & al. 2019; Soares & al. 2021).

A large part of the complex taxonomic history of Ingeae is centred around *Calliandra* Benth., the second largest genus of the tribe. *Calliandra* was first described by Bentham (1840) to accommodate 18 Neotropical species. Bentham (1844) later expanded *Calliandra* to include 60 species placed in five series, viz. *Calliandra* ser. *Macrophyllae* Benth., *C*. ser. *Laetevirentes* Benth., *C*. ser. *Pedicellatae* Benth., *C*. ser. *Nitidae* Benth. and *C*. ser. *Racemosae* Benth. In a later work, Bentham (1875) included four Asian species in *Calliandra*, viz. *C. cynometroides* Beddome, *C. geminata* Benth., *C. griffithii* Benth. and *C. umbrosa* Benth., thus extending the distribution of *Calliandra* into the Old World tropics. Two African species, *C. gilbertii* Thulin & Asfaw and *C. redacta* (J. H. Ross) Thulin & Hunde, were later included in the genus (Thulin & al. 1981). Additional species have been included in *Calliandra*, or newly described, over the years (e.g. Standley 1929; Barneby 1998).

In recent decades, *Calliandra* s.l. (sensu Bentham 1875 and subsequent work) has been split into several genera. The genus *Zapoteca* H. M. Hern. was described by Hernández (1986) to accommodate the approximately 25 species of *Calliandra* ser. *Laetevirentes* that he considered morphologically distinct from the other species of *Calliandra* s.l. Hernández (1986, 1989) argued that *Zapoteca* and *Calliandra* differ strikingly in seedling morphology, chromosome number (n=13 in *Zapoteca* vs. n=8 or 11 in *Calliandra*) and reproductive features. Moreover, species of *Zapoteca* have 16-grained acalymmate polyads (i.e. lacking a common exine) with circular thickenings on the pollen grains while the remaining species of *Calliandra* have 8-grained calymmate polyads (i.e. with a shared exine) and do not have any circular thickenings on the pollen grains. With further additions (Bässler 1998; Hernández 1989, 1990, 2015; Levin & Moran 1989; Hernández & Campos 1994; Hernández & Hanan-Alipi 1998), *Zapoteca* now consists of 23 species and 12 subspecies distributed in the Neotropics (and one species introduced to Africa and Asia; Hutchinson & Dalziel 1958, POWO 2021).

The most recent monograph of *Calliandra* s.l. (Barneby 1998) included approximately 130 New World species only, by implication thus making the genus strictly Neotropical. As a consequence, 18 species of *Calliandra* s.l. restricted to the Old World were left without any new or alternative generic placements. The Malagasy genus *Viguieranthus* Villiers was described by Villiers (2002) to accommodate eight of the species excluded from *Calliandra* s.l. by Barneby (1998). Villiers (2002) also described ten new species of *Viguieranthus*, raising the total number of species to 18. One species, *V. subauriculatus*, is, besides in Madagascar, also found in the Comoro Islands. Further, the genus *Afrocalliandra* E. R. Souza & L. P. Queiroz was established by Souza & al. (2013) to include the two African species described in *Calliandra* (*C. gilbertii* and *C. redacta*).

The genus *Thailentadopsis* Kosterm. was described by Kostermans (1977) to accommodate the single Thai species, *T. tenuis* (Craib) Kosterm., previously placed in *Pithecellobium* Mart. and *Acacia*. Kostermans (1977) argued that the new genus possessed a unique combination of morphological characters otherwise characteristic of several other mimosoid genera. Without making the new combinations, Nielsen (1981) referred *T. tenuis* to a broadly defined *Havardia* Small together with two other species, *Painteria nitida* (Vahl) Kosterm. from Sri Lanka and *Pithecellobium vietnamense* I. C. Nielsen from Vietnam. *Thailentadopsis* was resurrected by Lewis & Schrire (2003), including the above mentioned Asian species and adding *Calliandra geminata* (Wight & Arn.) Benth. to the synonymy of *T. nitida*. Furthermore, the Indian species *Calliandra cynometroides* Bedd. was excluded from *Calliandra* and placed in a monotypic genus, *Sanjappa* E. R. Souza & Krishnaraj, by Souza & al. (2016).

Two more taxa should be mentioned in association with *Calliandra* s.l.: the monotypic genera *Faidherbia* A. Chev. and *Guinetia* L. Rico & M. Sousa. *Faidherbia albida* (Delile) A. Chev. is distributed in Africa, Saudi Arabia, Jordan and Syria. It was originally described by Delile (1813) as *Acacia albida* Delile, but was later transferred to its own genus (Chevallier 1976) and included in Ingeae. *Guinetia* was established by Rico Arce & al. (1999) to accommodate the previously undescribed, Mexican species, *Guinetia tehuantepecensis* L. Rico & M. Sousa. However, Souza & al. (2013) transferred *Guinetia tehuantepecensis* to *Calliandra*. Subsequent phylogenetic studies assessed relationship of members of the ingoid clade, or parts thereof, but most studies focused mainly on *Acacia* and the (former) tribe Acacieae (e.g. Miller & al. 2003; Brown & al. 2008; Bouchenak-Khelladi & al. 2010; Miller & Seigler 2012), or on some of the alliances of the (former) tribe Ingeae (e.g., Iganci & al. 2016; Ferm & al. 2019; Koenen & al. 2020a), or had a broader focus in the Fabaceae or the mimosoid clade (e.g., Luckow & al. 2003; LPWG 2017; Koenen & al. 2020b), typically including few representatives of each included genus and sometimes with poor statistic support for many nodes.

Here, we investigate phylogenetic relationships in the ingoid clade (sensu Koenen & al. 2020a) using a substantially increased sample of taxa and/or data, in particular for the main focus of our study: relationships among the genera *Acaciella, Afrocalliandra, Calliandra* s.s., *Faidherbia, Sanjappa, Thailentadopsis, Viguieranthus* and *Zapoteca*. To date, only one phylogenetic study, that by the Legume Phylogeny Working Group (LPWG 2017), has included all these genera but the study was based on a single gene region (*matK*) and relationships were not resolved in the ingoid clade. We also include 27 additional genera representing the entire (pantropical) distribution of the ingoid clade, and we typically include multiple samples from each genus. The results are discussed in relation to gross morphology, geographical distribution, and existing classification of the studied species.

## Material and methods

### Data sampling

The phylogenetic tree presented by LPWG (2017) based on data from the plastid region *matK* was used as backbone for our taxon sampling. We selected 246 samples from the former tribe Ingeae and allied genera. Taxon names and authorities, voucher information and GenBank accessions for newly produced sequences are given in Appendix 1. Plant material of *Albizia* Durazz., *Calliandra* s.s., *Cojoba* Britton & Rose, *Enterolobium* Mart., *Inga* Mill., *Jupunba* Britton & Rose*, Pithecellobium, Samanea* (Benth.) Merr., *Senegalia, Vachellia* Wight & Arn.*, Zapoteca* and *Zygia* P. Browne was collected during field work in Ecuador, Jamaica and Puerto Rico 1995–2018. In addition, leaf material from species of *Viguieranthus* was obtained from the herbaria P and TAN, and material of *Cojoba, Enterolobium, Lysiloma* Benth. and *Pseudosamanea* Harms from the herbaria AAU, CICY, FTG and MO. In total, plant material from 54 samples not used in any previous phylogenetic study were used in the present study. Also, total DNA extracted by Ferm (2019) from 19 species of *Zapoteca* were used for DNA sequence amplification, as well as total DNA from *Z. portoricensis* (Jacq.) H. M. Hern. subsp. *portoricensis* and *Z. microcephala* (Britton & Killip) H. M. Hern., obtained from RBG Kew DNA Bank.

We selected five molecular markers for the present study: the plastid regions *matK, trnL-trnF* (including the *trnL* intron and the *trnL–trnF* spacer) and *ycf1*, and the nuclear ribosomal external transcribed spacer (ETS) and internal transcribed spacer (ITS). A total of 371 DNA sequences were newly produced for the present study, and analysed in combination with sequences downloaded from GenBank in order to get as complete datasets as possible. Four of the markers we selected (ETS, ITS, *matK* and *trnL-trnF*) have previously been used in phylogenetic studies of Fabaceae. In addition, we sequenced the entire plastid genome for three species (*Zapoteca* media [M. Martens & Galeotti] H. M. Hern., Z*. portoricensis* subsp. *portoricensis* and *Viguieranthus perrieri* [R. Vig] Villiers) in order to identify an informative region that has not been utilized in previous studies of the Fabaceae. The gene region *ycf1* showed the highest variation of all examined plastid DNA regions in these species, and we therefore chose to produce sequences of *ycf1* for the present study.

### DNA-extraction and whole genome sequencing

Total DNA was extracted and cleaned following the methods presented in Ferm & al. (2019). Total DNA was sent to the Science for Life Laboratory (SciLifeLab, Uppsala, Sweden) following the manufacturer’s instructions for the Illumina MiSeq platform (Illumina, San Diego, California, USA). Pair-end runs with 300-bp insert size fragments and 2 × 150 bp read lengths were performed. Library preparation was done using the Illumina SMARTer Thruplex DNAseq library preparation kit from Rubicon (Rubicon Genomics, Ann Arbor, Michigan, USA) at the SciLifeLab. The chloroplasts were assembled in Geneious Prime^®^ 2020.0.4 (https://www.geneious.com, Kearse & al. 2012) and all mapping was done using Geneious.

The raw data of *Zapoteca media* included 15798464 reads. The raw data of *Zapoteca portoricensis* subsp. *portoricensis* included 9553216 reads. The raw data of *Viguieranthus perrieri* included 12131060 reads. Reads of the raw data were paired with the insert size set to 350 bp using Geneious and trimmed using BBDuk v. 38.37 (by Brian Bushnell 2014), with the kmer lenght set to 27, reads < 35 bp to be discarded and a maximum lenght of reads set to 151 bp. Following trimming, 15590250 reads remained for *Z. media*, 9471114 reads remained for *Zapoteca portoricensis* subsp. *portoricensis* and 11999600 reads remained for *Viguieranthus perrieri*.

For *Z. media*, the paired reads were mapped to reference *Faidherbia albida* and the consensus sequence (conseq 1) extracted (367882 reads used). Next, the paired reads were mapped against conseq 1 and the consensus sequence extracted (conseq 2) (379173 reads used). The paired reads were mapped once again but against conseq 2 and the consensus sequence extracted (conseq 3) (407769 reads used). Used reads creating conseq 3 were assembled into contigs using De Novo assembly with SPAdes assembler v. 3.13.0 (http://cab.spbu.ru/software/spades/), and contigs > 500 bp were extracted. The contigs were mapped to reference *F. albida* and the consensus sequence extracted (conseq 4) (20 contigs used). Conseqs 1-4 were aligned to each other using Mauve (Darling & al. 2004) creating a final consensus sequence. The final consensus sequence was annotated using *F. albida* as reference.

For *Z. portoricensis* subsp. *portoricensis*, the paired reads were mapped to reference *Faidherbia albida* and the consensus sequence (conseq 1) extracted (365785 reads used). Next, the paired reads were mapped against conseq 1 and the consensus sequence extracted (conseq 2) (368071 reads used). Used reads creating conseq 2 were assembled into contigs using De Novo assembly with SPAdes assembler v. 3.13.0 (http://cab.spbu.ru/software/spades/), and contigs > 500 bp were extracted. The contigs were mapped to conseq 2 and the consensus sequence extracted (conseq 3) (19 contigs used). The contigs used for creating conseq 3 were mapped to the reference sequence *Faidherbia albida* and the consensus sequence extracted (conseq 4) (17 contigs used). Conseqs 1-4 were aligned to each other using Mauve (Darling & al. 2004) creating a final consensus sequence. The final consensus sequence was annotated using *F. albida* as reference.

For *V. perrieri*, the paired reads were mapped to reference *Faidherbia albida* and the consensus sequence (conseq 1) extracted (398957 reads used). Next, the paired reads were mapped against conseq 1 and the consensus sequence extracted (conseq 2) (404166 reads used). The paired reads were mapped once again but against conseq 2 and the consensus sequence extracted (conseq 3) (414377 reads used). Used reads creating conseq 3 were assembled into contigs using De Novo assembly with SPAdes assembler v. 3.13.0 (http://cab.spbu.ru/software/spades/), and contigs > 500 bp were extracted. The contigs were mapped to reference *F. albida* and the consensus sequence extracted (conseq 4) (20 contigs used). Conseqs 1-4 were aligned to each other using Mauve (Darling & al. 2004) creating a final consensus sequence. The final consensus sequence was annotated using *F. albida* as reference.

### Primer design and Sanger sequencing

The *ycf1* gene sequence was extracted from our amplified plastid genomes (of *Zapoteca media, Z. portoricensis* subsp. *portoricensis* and *Viguieranthus perrieri*, see above) and of four species downloaded from GenBank (*Pithecellobium flexicaule* [Benth.] J. M. Coult., *Inga leiocalycina* Benth., *Samanea saman* and *Faidherbia albida*), and were used as references for primer design conducted in Aliview v. 1.26 (Larsson 2014). The primer pairs were placed with a maximum of 1000 bp in between, contained a minimum of 40 percent CG-nucleotides and were 20-22 bp long. For *ycf1* 16 primers (combined into 12 different pairs) were designed. In addition, two new primer-pairs were designed for *matK*. The newly designed primers are listed in Table 1.

**Table 1.**
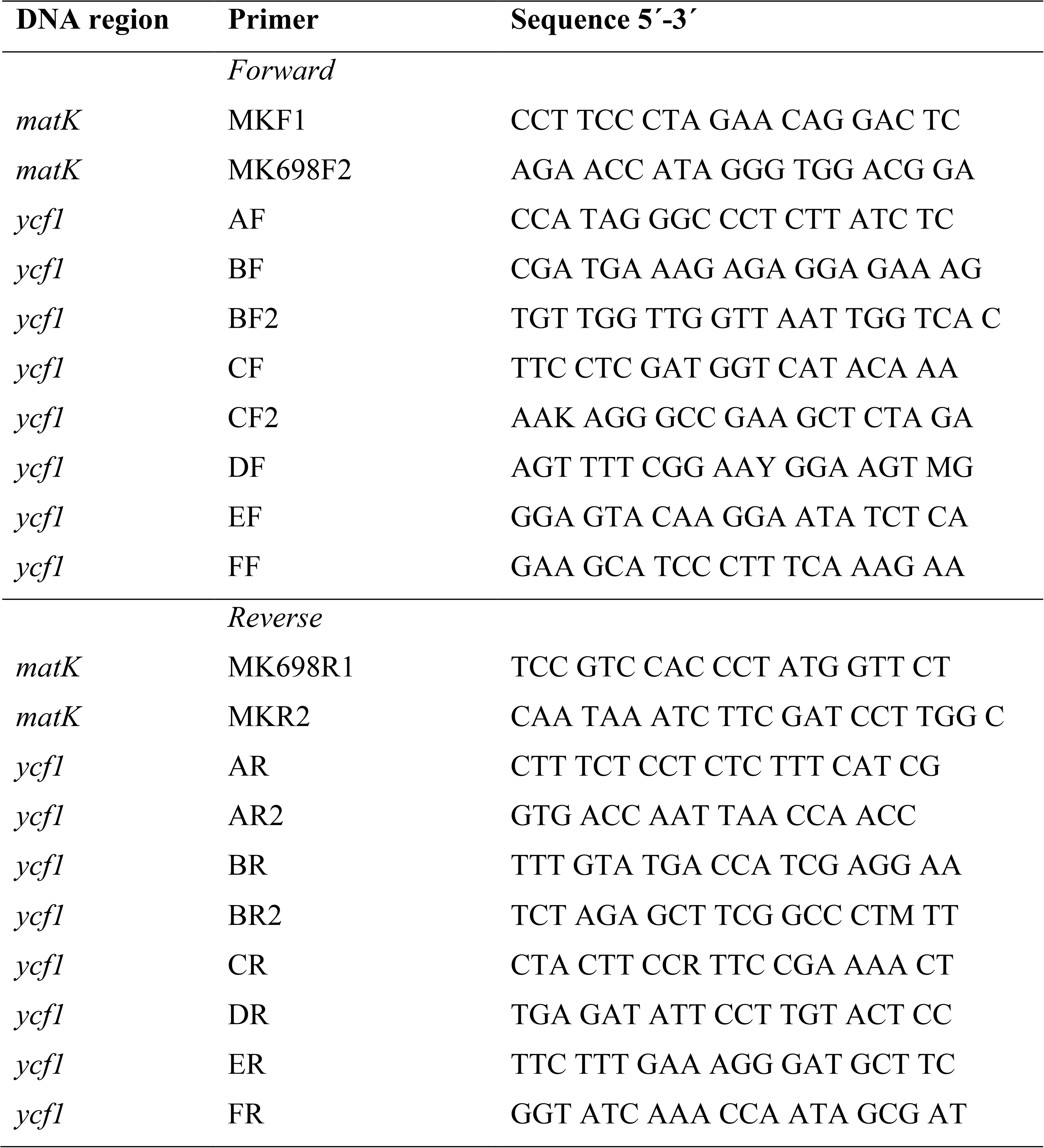
Primers newly designed for the present study.

For amplification of the five selected DNA regions for our sample of 103 specimens, PCR mixtures were prepared following standard protocols, as follows: 2,5 μL Paq5000 reaction buffer, 2 μL DNTP (containing 10 μM of each nucleotide), 0,3 μL of each primer, 0,25 μL BSA 1%, 2,5 μL TMACL, 0,25 μL Paq5000 DNA polymerase (5U/μL) and 1-3 μL DNA template. The amount of H2O added was adjusted between 13,9-15,9 μL in order to get a final sample volume of 25 μL. Nuclear ribosomal ETS was amplified using the primers 18S-IGS (Baldwin & Markos 1998) and AcR2 (Ariati & al. 2006) and amplifications were carried out as follows: 3-min initial denaturation at 95 °C followed by 35 cycles of 1-min denaturation at 95 °C, 1-min annealing at 55 °C and 1-min extension at 72 °C, and completed by a final extension of 7 min at 72 °C. Nuclear ribosomal ITS was amplified in two separate reactions using the primers P17 (Popp & Oxelman 2001) + ITS 491 (Ferm & al. 2019) and ITS 493 (Ferm & al. 2019) + 26S-82R (Popp & Oxelman 2001). Amplifications were carried out as follows: 1-min initial denaturation at 97 °C followed by 35 cycles of 10-s denaturation at 97 °C, 90-s annealing at 55 °C and 1-min extension at 72 °C, and completed by a final extension of 7 min at 72 °C. Plastid *matK* was amplified in two separate reactions using the primers MKF1 + MK698R1 and MK698F2 + MKR2 (all *matK* primers designed for this paper, see table 2). Amplifications were carried out as follows: 2-min initial denaturation at 95 °C followed by 35 cycles of 30-s denaturation at 95 °C, 30-s annealing at 52 °C (MK1F + MK698R1)/49 °C (MK698F2 + MK2R) and 1-min extension at 72 °C, and completed by a final extension of 7 min at 72 °C. Plastid *trnL-trnF* was amplified in two separate reactions using the primers c + d (Taberlet & al. 1991) and e (Taberlet & al. 1991) + jf1 (Ferm & al. 2019) and amplifications were carried out as follows: 3-min initial denaturation at 95 °C followed by 35 cycles of 1-min denaturation at 95 °C, 1-min annealing at 56 °C and 1-min extension at 72 °C, and completed by a final extension of 7 min at 72 °C. Plastid *ycf1* was amplified in separate reactions using primers in the following combinations: AF+AR; AF+AR2; BF+BR; BF+BR2; BF2+AR; BF2+BR2; CF+CR; CF2+BR; CF2+CR; DF+DR; EF+ER; FF+FR (all ycf1 primers designed for this paper, see table 2). Amplifications were carried out as follows: 3-min initial denaturation at 95 °C followed by 35 cycles of 1-min denaturation at 95 °C, 1-min annealing at 48 °C and 1-min extension at 72 °C, and completed by a final extension of 7 min at 72 °C. The PCR-products were purified using Illustra ExoProStar 1-Step (GE Healthcare) following the instructions from the manufacturer. Following purification, the samples were sent to Macrogen Europe in Amsterdam, the Netherlands, for sequencing. The same primers were used for sequencing as for PCR. Complementary strands of the obtained sequences were assembled and edited using Geneious Prime^®^ 2020.0.4 (https://www.geneious.com, Kearse & al. 2012).

### Phylogenetic analyses

Multiple alignments of the sequences were performed for each DNA region using MUSCLE (Edgar 2004) and adjusted by eye in AliView v.1.26 (Larsson 2014). The ETS and ITS datasets were concatenated into a combined nuclear dataset, and *matK, trnL*-*trnF* and *ycf1* were concatenated into a combined plastid dataset using Abioscripts (Larsson 2010). All trees were rooted on *Vachellia farnesiana* based on results in Kyalangalilwa & al. (2013) and LPWG (2017). The nuclear dataset and the plastid dataset were analysed separately and together, concatenated using Abioscripts (Larsson 2010), using maximum likelihood. Statistical support is here defined as having a aBayes support (aBS) of ≥ 0.95 or a bootstrap support (BS) of ≥ 95 (Alfaro & al. 2003; Erixon & al. 2003; Minh & al. 2013). Ultrafast bootstrap (UFBoot, a fast method for estimation of Bootstrap support under maximum likelihood; Minh & al. 2013; Hoang & al. 2018) was performed on IQ-TREE web server (Trifinopoulos & al. 2016; http://iqtree.cibiv.univie.ac.at) using IQ-TREE (Nguyen & al. 2015) with the default settings: automatic substitution model selection using ModelFinder including FreeRate heterogeneity (Kalyaanamoorthy & al. 2017), number of bootstrap alignments set to 1000, maximum iterations set to 1000 and minimum correlation coefficient set to 0.99 (Hoang & al. 2018); and with Approximate Bayes test (aBayes, a Bayesian-like transformation of aLRT; Anisimova & al. 2011) included. The maximum likelihood trees from the analyses were inspected in FigTree v.1.4.4 (Rambaut 2006-2016). The maximum likelihood tree resulting from the combined analysis was designed using Inkscape v.0.92 (https://inkscape.org). The aligned datasets from the separate analyses of the nuclear and plastid datasets are available in Dryad (https://datadryad.org/stash/share/kVGOkTngY2tFyjqL7pF3unNv2tK1q4_j-WI3S72CSAw).

## Results

### Single genome datasets

The results of the separate nuclear and plastid analyses are presented as supporting information (Suppl. Fig. 2-3). They show differences regarding relationships among genera but support is low for many nodes. Although statistic support is lacking, it is nevertheless worth noticing that the respective positions of a number of genera differ between results from nuclear vs. plastid data, e.g. the positions of *Acacia, Acaciella, Cojoba-Lysiloma, Faidherbia, Sanjappa* and *Thailentadopsis*.

### Combined nuclear and plastid dataset

The results of the analysis of the combined dataset (nuclear and plastid data; Fig. 2-3) are well-resolved. Generic relationships are generally well supported, while relationships within genera are sometimes less well statistically supported. The ingoid clade, i.e. all included taxa except *Vachellia*, is well-supported (aBayes support aBS 1/maximum likelihood bootstrap BS 100). Within this clade, *Acaciella* (aBS 1/BS 100) is sister to the remaining ingoid clade (aBS 1/BS 100). *Senegalia* (aBS 1/BS 100) is the next diverging clade, sister to remaining species (aBS1/BS 100). This clade (i.e. the traditional tribe Ingeae + the genus *Acacia*) comprises three major clades: clade 1 (aBS 1/BS 100) includes *Afrocalliandra* and *Calliandra* s.s.; clade 2 (aBS 1/BS 100) includes *Faidherbia, Sanjappa, Thailentadopsis, Viguieranthus* and *Zapoteca*; and clade 3 (aBS 1/BS 98) includes *Lysiloma, Cojoba, Havardia, Pithecellobium, Wallaceodendron* Koord., *Archidendron* F. Muell., *Archidendropsis* I. C. Nielsen, *Pararchidendron* I. C. Nielsen, *Albizia, Falcataria* (I. C. Nielsen) Barneby & J. W. Grimes, *Serianthes* Benth., *Cedrelinga* Ducke, *Paraserianthes* I. C. Nielsen, *Acacia, Punjuba* Britton & Rose, *Hydrochorea* Barneby & J. W. Grimes, *Jupunba, Balizia* Barneby & J. W. Grimes, *Pseudosamanea, Blanchetiodendron* Barneby & J. W. Grimes, *Leucochloron* Barneby & J. W. Grimes, *Enterolobium, Inga, Zygia, Chloroleucon* (Benth.) Record and *Samanea*. Clades 2 and 3 are sisters (aBS 1/BS100).

**Fig. 2.**
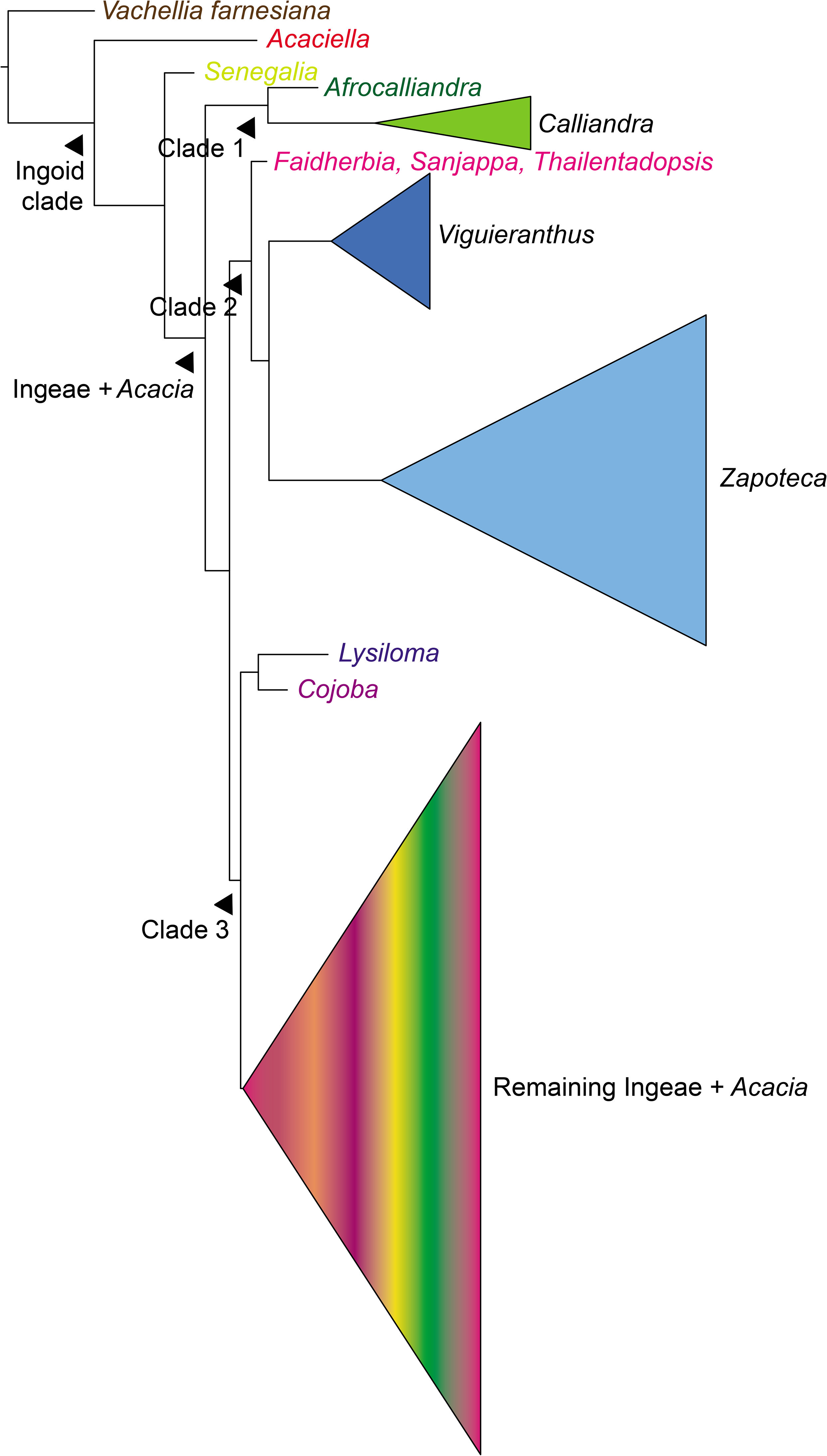
Phylogeny of the ingoid clade (Caesalpinioideae, Fabaceae), based on maximum likelihood analysis of the combined (nuclear and plastid) dataset (nuclear ribosomal ETS and ITS and plastid *matK, trnL-trnF* and *ycf1*), showing main results. The Ingeae and *Acacia* are resolved in three major clades indicated as clades 1–3.

**Fig. 3.**
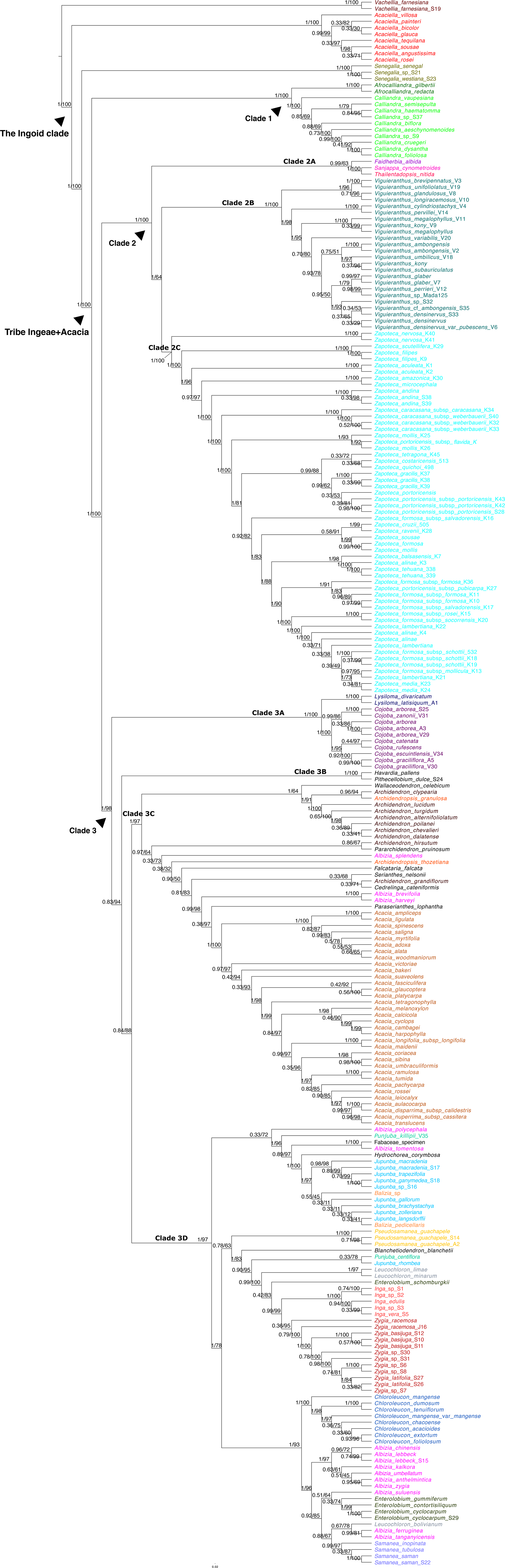
Phylogeny of the ingoid clade (Caesalpinioideae, Fabaceae) based on maximum likelihood analysis of the combined (nuclear and plastid) dataset (nuclear ribosomal ETS and ITS and plastid *matK, trnL-trnF* and *ycf1*). Support values at nodes are aBayes support values (aBS) and bootstrap support values (BS): presented aBS/BS. Strong support is defined as ≥ 0.95/95, according to recommendation in the literature.

Within clade 1, *Afrocalliandra* (aBS 1/BS 100) and *Calliandra* (aBS 1/BS 100) are each recovered as monophyletic and are found as sisters (aBS 1/BS 100). Within clade 2, three subclades are recovered. Subclade 2A (aBS 0.99/BS 63) consists of *Faidherbia* as sister to *Sanjappa* and *Thailentadopsis* (aBS 1/BS 100). Subclade 2B comprises *Viguieranthus* (aBS 1/BS 100), and subclade 2C *Zapoteca* (aBS 1/BS 100). Subclades 2B and 2C are sisters (aBS 1/BS 64). Within clade 3, four subclades are found. Subclade 3A (aBS 1/BS 100) comprises *Lysiloma* (aBS 1/BS 100) as sister to *Cojoba* (aBS 1/BS 100), and constitute the sister to remaining species (aBS 0.83/BS 94). Subclade 3B comprises *Pithecellobium* and *Havardia* (aBS 1/BS 100). Subclade 3C comprises *Wallaceodendron, Archidendron, Pararchidendron, Archidendropsis, Falcataria, Serianthes, Cedrelinga, Paraserianthes, Acacia* and some species of *Albizia*. Subclade 3D comprises *Punjuba, Hydrochorea, Jupunba, Balizia, Pseudosamanea, Blanchetiodendron, Leucochloron, Enterolobium, Inga, Zygia, Chloroleuchon, Samanea* and some species of *Albizia*.

### The position of *Acaciella*

The position of *Acaciella* differed among results; i.e., it was strongly supported as sister to the remaining ingoid clade based on nuclear ribosomal data (Suppl. Fig. 1) as well as in the combined analysis of plastid and nuclear ribosomal data (Fig. 3), whereas it was poorly supported as sister to the remaining ingoid clade except *Senegalia* based on plastid data only (Suppl. Fig. 2). We performed an additional total evidence analysis based on both nuclear ribosomal data and plastid data but with the single included ITS sequence of *Acaciella* removed. The result is yet another (unsupported) position of *Acaciella* as sister to clades 2 and 3 (Suppl. Fig. 3).

## Discussion

Several recent studies have pointed out the overall difficulties in resolving phylogenetic relationships within the Fabaceae (e.g. LPWG 2013, 2017; Koenen & al. 2020a, 2020b), something which is evident from the complex and instable classification of the Ingeae tribe and the genus *Calliandra* s.l. Our results show that the species of the former tribe Ingeae + *Acacia* is split into three major clades (Fig. 3; clades 1-3). *Acaciella* and *Senegalia* are resolved outside of these clades. The phylogenetic position of *Acaciella* has differed in previous work, and this is reflected also in our results. In the combined analyses and in analyses based on nuclear data, *Acaciella* is strongly supported as sister to all remaining species in the ingoid clade including *Sengalia* (Fig. 3, Suppl. Fig. 1). Although unsupported, the opposite is true for results based on plastid data, i.e. *Senegalia* is sister to the remaining species of the ingoid clade including *Acaciella* (Suppl. Fig. 2). Furthermore, in the combined analyses of nuclear and plastid data, with nuclear data excluded for *Acaciella*, the genus takes a third (unsupported) position, as sister to our clades 2 and 3 (Suppl. Fig 3). Taxon sampling can vary considerably between studies in the literature and may thereby prevent complete comparison, but previous studies based on plastid data (e.g., Bouchenak-Khelladi & al. 2010; Kyalangalilwa & al. 2013; LPWG 2017) have typically resolved *Acaciella* in a position congruent with our results from plastid data (Suppl. Fig. 1). However, in Miller & Seigler (2012), *Acaciella* and *Calliandra* are sisters.

Few phylogenetic studies of mimosoids have utilized nuclear data. Brown & al. (2008) resolved *Acaciella* as sister to the remaining ingoid clade based on analyses of nuclear ribosomal ETS and ITS, a result that is mimicked in our results based on nuclear data (Suppl. Fig. 1) as well as the combined analyses of nuclear and plastid data (Fig. 2-3). Miller & Seigler (2012) compared their results (*Acaciella* sister to *Calliandra*) with that of Brown & al. (2008) (*Acaciella* sister to remaining ingoid genera), and concluded that since their taxon sample was less dense than that of Brown & al. (2008), their detected sister-relationship between *Acaciella* and *Calliandra* may be an artefact caused by long-branch attraction (Miller & Seigler 2012). Our results would rather indicate cytonuclear discordance regarding the position of *Acaciella;* however, our results are ambiguous with no less than three alternative positions of *Acaciella* (Fig. 3 and Suppl. Fig. 1-3). Furthermore, results in Koenen & al. (2020a) indicate a sister-relationship between *Acaciella* and *Calliandra*, congruent with the results in Miller & Seigler (2012). However, this result in Koenen & al. (2020a) is not supported and they include only a single sample from each of these two genera. Despite considerable efforts, it thus seems clear that more research is needed regarding the deepest splits in the ingoid clade, employing a substantially increased sample of taxa and data from the nuclear and organellar genomes.

### Clade 1: *Afrocalliandra* and *Calliandra* s.s

The African genus *Afrocalliandra* (sensu Souza & al. 2013) and the Neotropical genus *Calliandra* s.s. (sensu Barneby 1998) are sisters (clade 1, Fig. 2-3) and together they represent the earliest diverging lineage within the Ingeae+*Acacia* clade (Fig. 2-3). Previous studies with a phylogenetic approach have resolved *Calliandra* s.s. as the sister to *Zapoteca* (Brown & al. 2008; Iganci & al. 2016; Ferm & al. 2019), but both genera were only represented by a few taxa in those studies, and no representatives of *Afrocalliandra, Viguieranthus, Thailentadopsis* or *Sanjappa* were included. An *Afrocalliandra-Calliandra* clade was detected in LPWG (2017) (excluding *Acaciella* with strong support), but its relationship to other ingoid taxa was not resolved. By contrast, Koenen & al. (2020a) found *Calliandra* as sister to *Acaciella*, but only a sample each from these genera were included and the result was not supported.

From a morphological perspective, *Afrocalliandra* and *Calliandra* s.s. are clearly distinguished from the other genera in the Ingeae+*Acacia* clade. Hernández proposed already in 1986 that *Calliandra* (sensu Hernández 1986) is unique within the Ingeae tribe (e.g. in polyad morphology), although the subsequent splits of *Calliandra* s.l. prevent exact comparison. Among features shared by *Afrocalliandra* and *Calliandra* s.s. are stigma- and cotyledon morphology (Thulin & al. 1981; Hernández 1986), and it has been argued that the species of *Afrocalliandra* should be included in *Calliandra* (Thulin & al. 1981). However, *Afrocalliandra* is restricted to Africa, *A. redacta* is found in South Africa and *A. gilbertii* in Kenya and Somalia, while *Calliandra* s.s. has a wide Neotropical distribution. Further, the two genera differ by having calymmate (*Calliandra* s.s.) vs. acalymmate (*Afrocalliandra*) polyads. *Calliandra* s.s. and *Afrocalliandra*, and several other ingoid taxa have the characteristic *“Calliandra-pod”*, but other genera that have this feature are readily distinguished from *Calliandra* s.s. and *Afrocalliandra* by having 16-grained, symmetrical, acalymmate polyads (Hernández 1986, 1989; Souza & al. 2016). By contrast, species of *Calliandra* s.s. have 8-grained, asymmetrical, calymmate polyads including a tail cell (Hernández 1986), and *Afrocalliandra* is variably reported to have 7-10 grains in their acalymmate polyads. According to Souza & al. (2013), *Afrocalliandra* has 7-grained, asymmetrical, acalymmate polyads including a tail cell. Uneven numbers of pollen grains in polyads are not uncommon within Ingeae (Barneby & Grimes 1997). However, Thulin & al. (1981) argue that *Afrocalliandra* has 8-grained polyads. Robbertse & von Teichman (1979) offer a possible solution to this apparent contradiction; they report that *A. redacta* has polyads with 7–10 pollen grains with 8-grained polyads being the most common condition. *Acaciella* has acalymmate, 8-grained polyads (Rico Arce & Bachman 2006), and is in that way similar to *Afrocalliandra*, but is native across the Neotropics and does not occur on the African continent.

Within *Calliandra* s.s. leaf morphology shows large variation between species and they can have one to many pairs of pinnae with one to many pairs of leaflets per pinna (Barneby 1998). The two species of *Afrocalliandra* have leaves with one pair of pinnae only, with several pairs of leaflets (Souza & al. 2013). Large variation of leaf morphology is also seen within *Acaciella*. The leaves have two to more than 25 pairs of pinnae and two to many pairs of leaflets per pinna (Rico Arce & Bachman 2006). *Acaciella, Calliandra* s.s. and *Afrocalliandra* all lack extra-floral nectaries (Marazzi & al. 2019). *Acaciella* and *Calliandra* s.s. do not possess spines or thorns (Hernández 1986; Barneby 1998; Rico Arce & Bachman 2006; Souza & al. 2013) with one exception, the Cuban *C. pauciflora*, which has stipules modified into spines (Barneby 1998). *Afrocalliandra gilbertii* has foliar stipules but thorns are sometimes formed from axillary branches while *A. redacta* has spiny stipules (Souza & al. 2013). Barneby (1998) stated that the stipular spines in *A. redacta* and in *C. pauciflora* probably are independently derived and thus not a sign of close evolutionary relationship since these two species differ in leaf morphology (*A. redacta* has spiral phyllotaxy and *C. pauciflora* has distichous phyllotaxy) and are well-separated geographically.

### The *“Calliandra* pod”

A specific fruit type, the “*Calliandra*-pod”, present in *Calliandra* s.s., *Afrocalliandra, Sanjappa, Viguieranthus* and *Zapoteca* has previously been mentioned as an indication of kinship within Ingeae (beginning with Bentham 1840). This type of pod has thickened margins and is dehiscent from the apex to the base with recurving valves. However, our results show that the “*Calliandra*-pod” is not a unique feature characteristic of a single clade; it occurs in clade 1 as well as in most members of clade 2. The “*Calliandra*-pod” has probably evolved independently several times within the Ingeae+*Acacia* clade. An alternative explanation for its occurrence in both clade 1 and clade 2 could be that the “*Calliandra*-pod” represents the ancestral state in the Ingeae+*Acacia* clade, with different pod types having evolved in *Faidherbia, Thailentadopsis* and in genera of clade 3. This seems, however, as a less likely scenario in the light of information in Barneby (1998), where he argues that a similar pod type as seen in *Calliandra* s.l. is found also in *Calliandropsis* H. M. Hern. & P. Guinet, some species of *Desmanthus* Willd., *Dichrostachys kirkii* Benth. and *Jacqueshuberia* Ducke. Since all of these taxa are positioned outside of the Ingeae+*Acacia* clade, *Jacqueshuberia* even outside of the mimosoid clade (LPWG 2017), it seems more probable that a similar fruit type, the so called “*Calliandra*-pod”, has evolved independently several times within the family of legumes.

### Clade 2: *Sanjappa, Thailentadopsis, Faidherbia, Viguieranthus* and *Zapoteca*

Within clade 2, the Old World species *Sanjappa, Thailentadopsis* and *Faidherbia* (clade 2A) form a group, which is sister to a clade comprising the Malagasy genus *Viguieranthus* and the Neotropical genus *Zapoteca* (Fig. 2-3; Suppl. Fig. 1 and 3). It should be noted, however, that plastid data result in another topology (Suppl. Fig. 2), where *Faidherbia* is sister to clades 2 and 3, and *Thailentadopsis* and *Sanjappa* are successive sisters to *Viguieranthus*. These results may tentatively indicate a reticulate (hybrid/allopolyploid) origin of these genera, but their positions in the plastid tree are not strongly supported and further studies (employing a dense taxon sample and data from the organellar and nuclear genomes) are needed to understand their evolutionary origins. *Faidherbia* has previously been suggested to be sister to *Zapoteca nervosa* (Urb.) H. M. Hern. (Miller & Seigler 2012), but our results refute a close relationship between *Faidherbia* and *Zapoteca*. In Souza & al. (2016), a *Sanjappa-Thailentadopsis-Faidherbia* clade was recovered, but as in the present study, they found cytonuclear discordance regarding their interrelationships (Souza & al. 2016: Fig. 1b). Further, the *Sanjappa*-*Thailentadopsis*-*Faidherbia* clade was sister to *Viguieranthus*, with *Zapoteca* excluded from this clade (Souza & al. 2016: Fig. 1a). This discrepancy compared to our results is, however, a rooting artefact, a consequence of the use of *Zapoteca* as outgroup in their study (see Souza & al. 2016, p. 2). Koenen & al. (2020a) found a clade comprising a single sample each of *Zapoteca, Viguieranthus*, and *Faidherbia*, a result that thus does not agree with our results where we use a denser taxon sampling. Complete comparison with our results is, however, complicated by the fact that no representatives of *Thailentadopsis* and *Sanjappa* were included in Koenen & al. (2020a).

### *Faidherbia, Sanjappa* and *Thailentadopsis*

*Faidherbia* differs from other ingoid species in some aspects, e.g. in having polyads with a variable number of pollen grains (16–32) (Kenrick & Knox 1982; Barnes & Fagg 2003) and in being deciduous during the rainy season (Barnes & Fagg 2003). By contrast, *Sanjappa* and *Thailentadopsis* share the feature of 16-grained acalymmate polyads with *Viguieranthus* and *Zapoteca* (Hernández 1986, 1989; Souza & al. 2016), a feature thus not consistently occurring in *Faidherbia*. Similarly, while *Sanjappa* has the same pod type as *Afrocalliandra, Calliandra* s.s., *Viguieranthus* and *Zapoteca* (Souza & al. 2016), different pod types are present in *Faidherbia* and *Thailentadopsis. Faidherbia* possesses an indehiscent coiled, twisted or falcate pod (Barnes & Fagg 2003). Coiled but otherwise dehiscent pods are also present in ingoid genera such as *Cojoba* and *Pithecellobium* (Barneby & Grimes 1996, 1997), none of which shows close relationship with *Faidherbia* in this study. *Thailentadopsis* has dehiscent, submoniliform pods (Lewis & Schirie 2003). Moniliform pods are also present in more distantly related ingoid taxa like *Albizia umbellata* (Lewis & Rico Arce 2005; POWO 2021), here positioned in clade 3 (Fig. 3). Further, *Faidherbia, Sanjappa*, and *Thailentadopsis* all have extra-floral nectaries, a feature also found in *Viguieranthus* but only in three species of *Zapoteca* (Villiers 2002; Barnes & Fagg 2003; Lewis & Schirie 2003; Souza & al. 2016) (see further below).

Regarding vegetative features, *Sanjappa* differs from almost all other ingoid taxa in having bifoliolate instead of bipinnate leaves, which is the most common feature in Ingeae (Souza & al. 2016). In that respect, *Sanjappa* resembles the species of *Inga*, a more distantly related genus here positioned in clade 3 (Fig. 3). Further, *Sanjappa, Thailentadopsis* and *Faidherbia* all possess spine-like stipules (Barnes & Fagg 2003; Lewis & Schirie 2003; Souza & al. 2016) while *Viguieranthus* and *Zapoteca* have leafy stipules. But, according to Villiers (2002), the stipules of *Viguieranthus* are “coriaceous to somewhat spiny”, but this is not mentioned for any specific species. In *Zapoteca, Z. aculeata* (Benth.) H. M. Hern. is the only species with spinescent stipules (Hernández 1989).

### *Viguieranthus* and *Zapoteca*

*Viguieranthus* and *Zapoteca* are here shown to be sisters (Fig. 2-3, Suppl. Fig. 1 and 3) (but see above on possible cytonuclear discordance regarding the positions of *Sanjappa, Thailentadopsis* and *Faidherbia*). Both *Viguieranthus* and *Zapoteca* have 16-grained acalymmate polyads and do not have spines or thorns, the only exception being *Z. aculeata* with stipular spines. Nevertheless, *Zapoteca* and *Viguieranthus* are clearly distinguished from each other, and are well separated geographically; *Viguieranthus* is endemic to Madagascar (with one species also occurring in the Comoro Islands) whereas *Zapoteca* has a Neotropical distribution (except for the recent introduction of *Z. portoricensis* to Africa and Asia; Hutchinson & Dalziel 1958; POWO 2021). The most common leaf structure in *Zapoteca* is to have more than one pair of pinnae with several pairs of leaflets/pinna, although a few species possess only one pair of pinnae and one to few pairs of leaflets (Hernández 1989). *Viguieranthus* on the other hand has leaves with one pair of pinnae only, with several pairs of leaflets/pinna that can be opposite or alternate in the same plant. Species of *Viguieranthus* have the stamens fused to each other and to the petals, and to the disk when present, forming a stamonozone (Villiers 2002), a feature not present in *Zapoteca*. In general, *Zapoteca* does not possess extra-floral nectaries, but there are three to four exceptions (viz. *Z. nervosa, Z. filipes* (Benth.) H. M. Hern and *Z. scutellifera* (Benth.) H. M. Hern., and occasionally also *Z. lambertiana* (G. Don) H. M. Hern., (Hernández 1989). By contrast, the species of *Viguieranthus* have a nectary on the apex on the petiole (Villiers 2002). Extra-floral nectaries are thus apparently ancestral in clade 2, since they are present in *Faidherbia, Sanjappa, Thailentadopsis, Viguieranthus* and in three early diverging species of *Zapoteca* (*Z. nervosa, Z. filipes* and *Z. scutellifera;* Fig. 3). It seems clear that the character must have been lost in the remaining species of *Zapoteca*, except for sometimes present in *Z. lambertiana*.

### Clade 3: remaining species of the Ingeae and *Acacia*

Within Clade 3, the Neotropical genera *Lysiloma* and *Cojoba* are sisters and this clade (clade 3A) is strongly supported as the sister to the remaining taxa in clade 3 (Fig. 2-3). This *“Cojoba* clade” is present in Koenen & al. (2020a), where it also includes the genus *Hesperalbizia* Barneby & J. W. Grimes. A “*Pithecellobium* clade” diverges next in Koenen & al. (2020a) and our results support that (clade 3B in Fig. 3). The remaining species of clade 3 comprise two sister clades (labelled clades 3C and 3D in Fig. 3). In clade 3C, the sister taxa *Wallaceodendron* and *Archidendron*, as well as one species of *Archidendropsis*, constitute the sister to a clade comprising *Acacia* and several genera for which our sampling is sparse (i.e., *Archidendropsis, Falcataria* and *Serianthes*), the monotypic genera *Pararchidendron, Paraserianthes* and *Cedrelinga*, and some species of *Archidendron* and *Albizia*. In clade 3D, two species of *Albizia*, as well as *Punjuba, Hydrochorea, Balizia* and most species of *Jupunba* are sister to two sister clades, of which one contains *Pseudosamanea, Blanchetiodendron*, one species each of *Punjuba* and *Jupunba, Inga, Zygia*, two species of *Leucochloron* and one species of *Enterolobium*. The second sister clade comprises *Chloroleucon*, most species of *Albizia and Enterolobium, Samanea* and one species of *Leucochloron*.

Results within clade 3 are well resolved but partly poorly supported. Our results show, however, that whereas *Acacia, Inga, Chloroleucon* and *Samanea* are monophyletic, the other included genera are not. Koenen & al. (2020a) focus more broadly of the mimosoids and include only a single representative of many genera, but some genera are shown to be non-monophyletic also in their study, i.e. *Albizia, Balizia*, and *Leucochloron*. Otherwise, our results may differ and most clades specified by Koenen & al. (2020a) in the equivialent of our clade 3 are not recovered in our results (i.e., the *Archidendron* clade, *Samanea* clade and *Albizia* clade of Koenen & al. 2020a). Earlier work is difficult to compare with; sampling in those studies may be too limited for the genera of our clade 3, and results may be poorly resolved.

Besides that the Neotropical genus *Chloroleucon* was recovered in a poorly supported unresolved clade in LPWG (2017) and represented by a single sample in Koenen & al. (2020a), the phylogenetic position of *Chloroleucon* has not previously been investigated using molecular data. It (i.e., *Chloroleucon tenuiflorum* (Benth.) Barneby & J. W. Grimes) was sister to *Samanea saman* (Jacq.) Merr. in Koenen & al. (2020a), but their proposed “*Samanea* clade” is not present in our results. Instead, we find that *Chloroleucon* is well supported as sister to a larger clade also comprising taxa excluded from the *“Samanea* clade” of Koenen & al. (2020a), i.e., some species of *Albizia*, most species of *Enterolobium*, and one species of *Leucocloron* (Fig. 3). *Chloroleucon* is characterized by having axillary thorns, striate resting buds and by flowering before the yearly production of new leaves. Together with *Leucochloron, Blanchetiodendron, Cathormion* (now included in *Albizia*; Koenen & al. 2020a) and *Thailentadopsis*, it was placed in the “*Chloroleucon* alliance” by Lewis & Rico Arce (2005), but this informal group does not represent a monophyletic group in our results. while all genera of the *“Chloroleucon* alliance” of Lewis & Rico Arce (2005) (except *Thailentadopsis*) are found in our clade 3, they do not form a clade. *Thailentadopsis* is found in clade 2 (Fig. 3; clade 2A).

In our results *Leucochloron* is not confirmed as monophyletic. *Leucochloron limae* Barneby & J. W. Grimes and *L. minarum* (Harms) Barneby & J. W. Grimes are strongly supported as sisters and found in a strongly supported clade together with *Inga, Zygia* and *Enterolobium schomburgkii* Benth. (Fig 1; clade 3C). *Leucochloron bolivianum* C. E. Hughes & Atahuachi is found in an unresolved position within clade 3D. The four remaining species of *Enterolobium* included in this study are found in a clade that is poorly supported (Fig. 3; clade 3D). *Leucochloron, Inga* and *Zygia* have previously been shown to be closely related, also including *Macrosamanea* Britton & Rose (Ferm & al. 2019), *Abarema cochliacarpos* (Gomes) Barneby & J. W. Grimes (LPWG 2017), and *Blanchetiodendron* (Koenen & al. 2020a) in the same clade. However, the *“Inga* alliance” defined by Lewis & Rico Arce (2005), including *Macrosamanea, Zygia* and *Inga* as well as other genera, i.e. *Guinetia* (now included in *Calliandra* s.s.), *Calliandra* (sensu Barneby 1998), *Viguieranthus, Cojoba, Cedrelinga* and *Archidendron*, do not comprise a monophyletic group. Since *Leucochloron limae, L. minarum* and *Enterolobium schomburgkii* are here shown to be more closely related to *Inga* and *Zygia* than to the other species of these respective genera. Similar results were indicated in LPWG (2017), but *Inga* is characterized by having pinnate leaves and *Zygia* by having cauli-and/or ramiflory, neither of which apply to *L. limae, L. minarum* or *Enterolobium schomburgkii*. Moreover, these two genera are clearly not monophyletic (Fig. 3), and we argue that the status of *Leucochloron* and *Enterolobium* need to be further evaluated and probably revised.

## Conclusions

Despite considerable efforts, relationships among mimosoids are still not fully understood. Relationships presented in the literature are often inconsistent and/or poorly resolved and supported, and the reasons for these problems need more research. Many studies have had a broad focus within the mimosoids or the entire legume family, and sampling of taxa may therefore have been sparse for individual genera. This is particularly true regarding the main focus of our study, the genera *Acaciella, Afrocalliandra, Calliandra* s.s., *Faidherbia, Sanjappa, Thailentadopsis, Viguieranthus* and *Zapoteca*. Evidence of cytonuclear discord, conflicting topologies retrived based on data from the organellar genome(s) vs. the nuclear genome, seen in our results as well as when comparing our results with those of other studies, may have different reasons, including methodological factors such as sampling error(s) and suboptimal model selection as well as biological factors including hybridization/introgression and incomplete lineage sorting. Biological reasons cannot be ruled out at this point, and it is possible that future studies utilizing genomic data from several genomes and a dense sample of taxa may provide clarity.

Based on our results, the ingoid clade (sensu Koenen & al. 2020a) comprises three major clades, clade 1 (*Afrocalliandra* and *Calliandra*), clade 2 (a *Faidherbia – Sanjappa – Thailentadopsis* clade sister to a *Vigueranthus – Zapoteca* clade), and clade 3 (*Lysiloma, Cojoba, Havardia, Pithecellobium, Wallaceodendron, Archidendron, Archidendropsis, Pararchidendron, Albizia, Falcataria, Serianthes, Cedrelinga, Paraserianthes, Acacia, Punjuba, Hydrochorea, Jupunba, Balizia, Pseudosamanea, Blanchetiodendron, Leucochloron, Enterolobium, Inga, Zygia, Chloroleucon* and *Samanea*). Outside of these three major clades are the genera *Acaciella* and *Senegalia*, and the position of the former is unclear with weak (unsupported) evidence of cytonuclear discordande detected here as well as by deviating phylogenetic results in the literature. The three major clades and subclades within them are typically difficult to characterize morphologically. However, the traditionally described *“Calliandra-pod”* is a misconception, at least from an evolutionary perspective. It is not only found in the taxa formerly referred to *Calliandra* (i.e. *Calliandra* s.l.), but also in other legume genera. *Calliandra* s.s. is the only genus within the Ingeae + *Acacia* clade with calymmate polyads, which makes it easy to distinguish from the other genera of this clade.

Finally a few words about the geographic distribution of the ingoid clade, which is pantropical with most of its diversity in the Neotropics. The crown age of the Ingeae + *Acacia* clade has been estimated at 23.9±3.1 myr (Lavin & al. 2005) and, consequently, the distribution of the clade seen today cannot have been caused by plate tectonics. Instead, long distance dispersal, e.g. across the Atlantic (and other oceans to reach Australia), has apparently occurred several times in the clade. Our clade 1 includes the African *Afrocalliandra* sister to the Neotropical *Calliandra* s.s. Clade 2 includes the Old World genera *Faidherbia, Sanjappa, Thailentadopsis* and *Viguieranthus*, as well as the Neotropical *Zapoteca*. Clade 3 includes both New World and Old World genera, along with species from Australia, an area which is not represented in clades 1 and 2. Given that the Ingeae + *Acacia* clade has its origin in the Old World, at least three dispersal events to the New World must have occurred, once in clade 1 and once in clade 2, and at least once in clade 3. The most likely way of dispersal of pods and seeds across the Atlantic is by means of oceanic surface currents (Töpke & Song 2020) from Africa to South America (Renner 2004). Seeds of *Zapoteca* have been shown to be viable even after up to 70 days in saline water (Hernández 1989). Thus, germination of *Zapoteca* and probably other ingoid seeds would be possible even after a long ocean travel.

## Acknowledgements

We thank the curators and the staff at AAU, CICY, FTG, MO and P for allowing access to material for DNA-extractions. We also thank the curators and the staff of TAN for providing material for DNA-extraction and Dr. Sylvain Razafimandimbison (The Swedish Museum of Natural History) for bringing the samples. We thank Dr. Rodrigo Duno de Stefano (Centro de Investigación Científica de Yucatán) for sharing collected leaf material for DNA-extractions and Steve Maldonado Silvestrini for supplying newly collected leaf material. The first author, JF, thanks Dr. James Ackerman and Dr. Frank Axelrod (University of Puerto Rico Rio Piedras) for assistance during field work in Puerto Rico. We thank Dr. Alvaro Pérez at Universidad Católica, Quito, for obtaining collecting permits in Ecuador (no: 1-2016-IC-FL0-DNB/MA) including BS and JF. We also thank National Environment and Planning Agency in Jamaica for issuing a collecting and exportation permit for JF (no: 18/27) and the herbarium at University of Puerto Rico Rio Piedras for providing permission to collect and export plants in Puerto Rico. The study was supported by funds from Stiftelsen Lars Hiertas minne and Helge Ax:son Johnsons stiftelse to JF, and from the Regnell foundation, Uppsala University to BS, and from the Royal Swedish Academy of Sciences and Stockholm University to CR.

## Supplementary files captions

Suppl. Fig. 1. Maximum likelihood analysis of the nuclear dataset (ETS and ITS). Values at nodes are aBayes support values (aBS) and bootstrap support values (BS): presented aBS/BS.

Suppl. Fig. 2. Maximum likelihood analysis of the plastid dataset (*matK, trnL-trnF* and *ycf1*). Values at nodes are aBayes support values (aBS) and bootstrap support values (BS): presented aBS/BS.

Suppl. Fig. 3. Maximum likelihood analysis of the combined dataset (nuclear and plastid), with *Acaciella* represented by plastid sequences only. Values at nodes are aBayes support values (aBS) and bootstrap support values (BS): presented aBS/BS.

## Appendix 1. Taxon names and GenBank accession numbers of DNA sequences included in this study

Voucher data is given for accessions for which DNA sequences were newly obtained, using the following format: Taxon name, country, collector and collector number, herbarium code, GenBank accession numbers (ETS, ITS, *matK, trnL-trnF, ycf1*), ycf1 sequences not available in GenBank but in Dryad. – missing data; * newly generated sequence.

***Acaciella** angustissima* (Mill.) Britton & Rose; EF638082.1, EF638169.1, EU812043.1, HM020825.1, –; *Acaciella bicolor* Britton & Rose; –, –, –, HM020826.1, –; *Acaciella glauca* (L.) L. Rico; –, –, EU812042.1, DQ371857.1, –; *Acaciella painteri* Britton & Rose; –, –, –, HM020828.1, –; *Acaciella rosei* (Standl.) Britton & Rose; –, –, –, HM020829.1, –; *Acaciella sousae* (L. Rico) L. Rico; –, –, –, HM020830.1, –; *Acaciella tequilana* (S. Watson) Britton & Rose; –, –, EU812044.1, HM020831.1, –; *Acaciella villosa* (Sw.) Britton & Rose; –, –, KX302289.1, –, –; ***Afrocalliandra** gilbertii* (Thulin & Hunde) E. R. Souza & L. P. Queiroz; –, JX870690.1, KX581218.1, –, –; *Afrocalliandra redacta* (J. H. Ross) E. R. Souza & L. P. Queiroz; –, JX870732.1, KX581219.1, JX870853.1, –; ***Acacia** adoxa* Pedley; EF638087.1, AF360715.1, AF523076.1, JF420480.1, –; *Acacia alata* R. Br.; EF638089.1, AF360699.1, JF420001.1, JF420541.1, –; *Acacia ampliceps* Maslin; EF638117.1, KC200598.1, AF523074.1, AF522983.1, –; *Acacia aulacocarpa* A. Cunn ex Benth.; JF420289.1, JF420068.1, AF274214.1, JF420501.1, –; *Acacia bakeri* Maiden; –, –, KM894854.1, KC957765.1, –; *Acacia calcicola* Forde & Ising; JN935146.1, KC200685.1, AF274220.1, KC957695.1, –; *Acacia cambagei* R. T. Baker; –, –, JX850060.1, –, –; *Acacia coriacea* DC.; KC283745.1, KC200735.1, AY180923.1, KC957704.1, –; *Acacia cyclops* F. Muell.; KT149821.1, JF420024.1, JQ412187.1, JF420460.1, –; *Acacia disparrima* subsp. *calidestris* M. W. McDonald & Maslin; KC283548.1, KC200792.1, KC421478.1, KC958353.1, –; *Acacia fasciculifera* F. Muell. ex Benth.; –, AF487769.1, AF274154.1, –, –; *Acacia glaucoptera* Benth.; KC283430.1, KC200841.1, AF274217.1, KC958175.1, –; *Acacia harpophylla* F. Muell. ex Benth.; JN935163.1, KC200851.1, KM894642.1, KC958015.1, –; *Acacia leiocalyx* (Domin) Pedley; KC283381.1, KC200908.1, AF274216.1, KC958123.1, –; *Acacia ligulata* Aiton ex Steud.; KR994976.1, KC200918.1, AF274155.1, JF420464.1, –; *Acacia longifolia* (Andrews) Willd. subsp. *longifolia;* HM007633.1, HM007658.1, KC421317.1, KC958105.1, –; *Acacia maidenii* F. Muell.; KT149845.1, HM007664.1, KM894773.1, –, –; *Acacia melanoxylon* R. Br.; KT149850.1, JF420093.1, AF274166.1, KJ782136.1, –; *Acacia myrtifolia* Willd.; JN935177.1, FJ868400.1, AF274160.1, –, –; *Acacia nuperrima* subsp. *cassitera* Pedley; KC283404.1, KC200968.1, –, KC958154.1, –; *Acacia pachycarpa* F.Muell. ex Benth.; FJ868443.1, KC200991.1, AF274153.1, –, –; *Acacia platycarpa* F. Muell.; DQ029285.1, DQ029244.1, AF274223.1, KC958041.1, –; *Acacia ramulosa* W. Fitzg.; KC283833.1, KC201034.1, KC013731.1, JX870552.1, –; *Acacia rossei* Standl.; FJ868413.1, AF487756.1, AF274162.1, –, –; *Acacia saligna* (Labill.) H. L. Wendl.; KM095744.1, JF420109.1, HM020727.1, HM020817.1, –; *Acacia sibina* Maslin; JN935190.1, –, KC013746.1, JX870567.1, –; *Acacia spinescens* Benth.; EF638090.1, AF360700.1, AF523082.1, AF195706.1, –; *Acacia suaveolens* Willd.; FJ868451.1, KC201106.1, AF274221.1, JF420482.1, –; *Acacia tetragonophylla* F. Muell.; KT149865.1, KC201122.1, KC013726.1, KC958066.1, –; *Acacia translucens* Hook.; EF638112.1, AF360722.1, AF274165.1, AF522984.1, –; *Acacia tumida* F. Muell. ex Benth.; EF638114.1, AF360709.1, AF523111.1, AF522986.1, –; *Acacia umbraculiformis* Maslin & Buscumb; KC283593.1, KC201147.1, KC013757.1, JX870578.1, –; *Acacia victoriae* Benth.; DQ029322.1, DQ029281.1, AF274226.1, –, –; *Acacia woodmaniorum* Maslin & Buscumb; KC283618.1, –, KC013854.1, JX870600.1, –; ***Albizia** anthelmintica* Baill.; –, MN257768.1, KX302295.1, –, –; *Albizia brevifolia* Schinz; –, –, KX302300.1, –, –; *Albizia chinensis* (Osbeck.) Merr.; –, KP092696.1, LM643809.1, LM643812.1, –; *Albizia ferruginea* Benth.; –, –, KX302303.1, –, –; *Albizia harveyi* Fourn.; –, –, AF523075.1, EU439977.1, –; *Albizia kalkora* (Roxb.) Prain; EF638158.1, MH710962.1, HQ427295.1, AF522945.1, –; *Albizia lebbeck* (L.) Benth.; EF638155.1, KX057828.1, EU812047.1, EU440023.1, –; (S15) Puerto Rico, Ferm 106 (UPRRP); MW849561*, MZ015524*, MZ169674*, MW940450*, *ycf1*\*; *Albizia polycephala* (Benth.) Killip; KF921625.1, KF933275.1, –, –, –; *Albizia splendens* Miq.; –, –, KX302312.1, –, –; *Albizia suluensis* Gerstner; –, –, JX517858.1, –, –; *Albizia tanganyicensis* Baker f.; –, –, JF270636.1, –, –; *Albizia tomentosa* Standl.; –, –, AF523093.1, AF522994.1, –; *Albizia umbellatum* (Vahl) Kosterm.; EF638157.1, EF638182.1, AF274122.1, AF522949.1, –; *Albizia zygia* J. F. Macbr.; –, KX057829.1, KX302313.1, KX268144.1, –; ***Archidendron** alternifoliolatum* (T. L. Wu) I. C. Nielsen; –, KR531966.1, KR530670.1, –, –; *Archidendron chevalieri* (Kosterm.) I. C. Nielsen; –, –, LC080890.1, –, –; *Archidendron clypearia* (Jack) I. C. Nielsen; –, KP092698.1, KJ510955.1, –, –; *Archidendron dalatense* (Kosterm.) I. C. Nielsen; –, KR531776.1, KR530397.1, –, –; *Archidendron grandiflorum* (Soland. ex Benth.) I. C. Nielsen; –, –, KM894775.1, –, –; *Archidendron hirsutum* I. C. Nielsen; –, –, EU361860.1, AF365042.1, –; *Archidendron lucidum* (Benth.) I. C. Nielsen; –, KT321363.1, HQ415282.1, –, –; *Archidendron poilanei* (Kosterm.) I. C. Nielsen; –, –, LC080888.1, –, –; *Archidendron turgidum* (Merr.) I. C. Nielsen; –, KP092711.1, KP094140.1, –, –; ***Archidendropsis** granulosa* (Labill.) I. C. Nielsen; –, –, KX302316.1, –, –; *Archidendropsis thozetiana* (F. Muell.) I. C. Nielsen; –, EF638179.1, KM894536.1, –, –; ***Balizia** pedicellaris* (DC.) Barneby & J. W. Grimes; –, JX870657.1, KX302318.1, JX870789.1, –; *Balizia* Barneby & J. W. Grimes sp.; –, –, KX302319.1, –, –; ***Blanchetiodendron** blanchetii* (Benth.) Barneby & J. W. Grimes; KF921626.1, JX870658.1, KX302320.1, JX870790.1, –; ***Calliandra** aeschynomenoides* Benth.; –, JX870659.1, –, JX870791.1, –; *Calliandra biflora* Tharp; –, JX870667.1, –, JX870798.1, –; *Calliandra cruegeri* Griseb.; –, JX870679.1, –, JX870808.1, –; *Calliandra dysantha* Benth.; EF638121.1, JX870684.1, –, JX870813.1, –; *Calliandra foliolosa* Benth.; EF638122.1, EF638181.1, MG718924.1, –, –; *Calliandra haematomma* (DC.) Benth.; –, JX870695.1, –, JX870822.1, –; *Calliandra semisepulta* Barneby; –, JX870737.1, –, JX870856.1, –; *Calliandra vaupesiana* R. S. Cowan in R. E. Schult.; –, JX870754.1, KR270507.1, JX870870.1, –; *Calliandra* Benth. sp. (S9) Ecuador, Ferm 10 (Q); MW849572*, MZ015525*, –, MW940452*, *ycf1*\*; (S37) Jamaica, Ferm 100 (IJ); MW849571*, –, MZ169635*, MW940451*, *ycf1*\*; ***Cedrelinga** cateniformis* (Ducke) Ducke; –, JX870757.1, KX302323.1, JX870873.1, –; ***Chloroleucon** acacioides* (Ducke) Barneby & J. W. Grimes; KF921629.1, KF921672.1, –, –, –; *Chloroleucon chacoense* (Burkart) Barneby & J. W. Grimes; –, –, KY046033.1, –, –; *Chloroleucon dumosum* (Benth.) G. P. Lewis; KF921632.1, KF921680.1, KX581225.1, KF921756.1, –; *Chloroleucon extortum* Barneby & J. W. Grimes; KF921636.1, KF921681.1, KX581226.1, KF921760.1, –; *Chloroleucon foliolosum* (Benth.) G. P. Lewis; KF921640.1, KF921686.1, –, –, –; *Chloroleucon mangense* Britton & Rose; –, –, AY386921.1, AF522950.1, –; *Chloroleucon mangense* var. *mangense;* KF921645.1, KF921690.1, –, –, –; *Chloroleucon tenuiflorum* (Benth.) Barneby & J. W. Grimes; KF921646.1, KF921691.1, –, –, –; ***Cojoba** arborea* (L.) Britton & Rose; EF638095.1, EF638186.1, KX302324.1, JX870874.1, –; (A3) Mexico, R. Duno 2348 (CICY) MW849552*, MZ015526*, –, MW940453*, *ycf1*\*; (V29) Mexico, P. Tenorio L. 489 (AAU) MW849554*, MZ015528*, –, –, –; (S25) Puerto Rico, Ferm 116 (UPRRP) MW849553*, MZ015527*, –, MW940454*, *ycf1*\*; *Cojoba catenata* (Donn. Sm.) Britton & Rose; –, –, AY944554.1, AY944538.1, –; *Cojoba escuintlensis* (Lundell) L. Rico (V34) Mexico, F. Ventura A. 20002 (AAU) MW849555*, MZ015529*, MZ169641*, –, *ycf1*\*; *Cojoba graciliflora* (S. F. Blake) Britton & Rose (A5) Mexico, R. Duno 2550 (CICY) MW849556*, MZ015530*, –, –, *ycf1*\*; (V30) Mexico, F. Ventura A. 20849 (AAU) MW849557*, MZ015531*, MZ169636*, MW940455*, *ycf1*\*; *Cojoba rufescens* Britton & Rose; –, –, GQ981971.1, –, –; *Cojoba zanonii* (Barneby) Barneby & J. W. Grimes (V31) Dominican Republic, Jestrow 2012-291 with T. Clase, C. Husby & J. Lopez (FTG) MW849558*, MZ015532*, MZ169628*, MW940456*, *ycf1*\*; ***Enterolobium** contortisiliquum* (Vell.) Morong; EF638151.1, EF638190.1, AF274124.1, AF522952.1, –; *Enterolobium cyclocarpum* (Jacq.) Griseb.; EF638150.1, EF638191.1, AY650277.1, AF522953.1, –; (S29) Puerto Rico, Ferm 108 (UPRRP); MW849559*, MZ015533*, –, MW940457*, –; *Enterolobium gummiferum* J. F. Macbr.; KF921652.1, KF921696.1, KX581227.1, KF921773.1, –; *Enterolobium schomburgkii* Benth.; KF921653.1, KF921697.1, GQ981984.1, KF921774.1, –; **Fabaceae** specimen; –, –, HM020734.1, HM020827.1, –; ***Faidherbia** albida* (Delile) A. Chev.; EF638163.1, KX057872.1, HM020737.1, KR997877.1, –; ***Falcataria** moluccana* (Miq.) Barneby & J. W. Grimes; HM800430.1, MG751361.1, KX458480.1, –, –; ***Havardia** pallens* Britton & Rose; KF921656.1, KF921698.1, AF274125.1, AF522955.1, –; ***Hydrochorea** corymbosa* (Rich.) Barneby & J. W. Grimes; KF921657.1, JX870763.1, KX302331.1, JX870879.1, –; ***Inga** edulis* Mart.; KF921658.1, JX870764.1, AF523078.1, JX870880.1, –; *Inga vera* Willd. (S5) Puerto Rico, Ferm 119 (UPRRP); MW849550*, –, –, MW940462*, –; *Inga leiocalycina* Benth.; KT428296.1; *Inga* Mill. sp (S1) Ecuador, Ferm 32 (Q); MW849547*, MZ015535*, –, MW940459*, *ycf1*\*; (S2) Ecuador, Ferm 30 (Q); MW849548*, –, –, MW940460*, *ycf1*\*; (S3) Ecuador, Ferm 57 (Q); MW849549*, –, –, MW940461*, *ycf1*\*; ***Jupunba** brachystachya* (DC.) M. V. B. Soares, M. P. Morim & Iganci; –, –, KX374503.1, –, –; *Jupunba gallorum* (Barneby & J. W. Grimes) M. V. B. Soares, M. P. Morim & Iganci; –, –, KX374508.1, –, –; *Jupunba ganymedea* (Barneby & J. W. Grimes) M. V. B. Soares, M. P. Morim & Iganci (S18) Ecuador, Ferm 52 (Q); MW849566*, –, –, MW940447*, *ycf1*\*; *Jupunba trapezifolia* (Vahl.) Moldenke; EF638110.1, EF638166.1, –, –, –; *Jupunba langsdorffii* (Benth.) M. V. B. Soares, M. P. Morim & Iganci; –, –, KX374510.1, –, –; *Jupunba rhombea* (Benth.) M. V. B. Soares, M. P. Morim & Iganci; –, –, KX374512.1, –, –; *Jupunba macradenia* (Pittier) M. V. B. Soares, M. P. Morim & Iganci; –, –, KX374513.1, –, –; (S17) Ecuador, Ferm 51 (Q); MW849573*, MZ015522*, MZ169632*, MW940448 *, *ycf1*\*; *Jupunba zolleriana* (Standl. & Steyerm.) M. V. B. Soares, M. P. Morim & Iganci; –, –, KX374515.1, –, –; *Jupunba* Britton & Rose sp. (S16) Ecuador, Ferm 26 (Q); MW849574*, MZ015523*, MZ169633*, –,*ycf1*\*; ***Leucochloron** bolivianum* C. E. Hughes & Atahuachi; KF921660.1, KF921699.1, KX581230.1, KF921776.1, –; *Leucochloron limae* Barneby & J. W. Grimes; KF921663.1, JX870766.1, KX302334.1, JX870882.1, –; *Leucochloron minarum* (Harms) Barneby & J. W. Grimes; KF921664.1, KF921702.1, KX581231.1, KF921779.1, –; ***Lysiloma** divaricatum* Benth.; EF638093.1, AF487755.1, AF523088.1, HM020837.1, –; *Lysiloma latisiliquum* (L.) Benth. (A1) Mexico, Sima et al. 2287 (CICY) MW849567*, MZ015536*, MZ169642*, –, *ycf1*\*; ***Pararchidendron** pruinosum* (Benth.) I. C. Nielsen; EF638129.1, JF420082.1, AF274127.1, EU439985.1, –; ***Paraserianthes** lophantha* (Willd.) I. C. Nielsen KU727943.1, EF638204.1, AF274128.1, AF522962.1, –; ***Pithecellobium** dulce* (Roxb.) Benth. (S24) Puerto Rico, Ferm 107 (UPRRP); MW849560*, MZ015540*, MZ169638*, MW940466*, *ycf1*\*; *Pithecellobium flexicaule* (Benth.) J. M. Coult.; KX852444.1; ***Pseudosamanea** guachapele* (Kunth) Harms; KF921667.1, JX870769.1, AF523079.1, AF522964.1, –; (S14) Ecuador, Ferm 40 (Q); MW849563*, –, –, MW940449*, –; (A2) Mexico, Reyes-Garcia & Gomez 4511 (MO) MW849562*, MZ015541*, MZ169623*, –, *ycf1*\*; ***Punjuba** centiflora* (Barneby & J. W. Grimes) M. V. B. Soares, M. P. Morim & Iganci; –, –, KX374504.1, –, –; *Punjuba killipii* Britton & Rose ex Britton & Killip (V35) Ecuador, Homeier 359 (MO); MW849565*, MZ015542*, –, MW940467*, *ycf1*\*; ***Samanea** inopinata* (Harms) Barneby & J. W. Grimes; –, –, KX581234.1, –, –; *Samanea saman* (Jacq.) Merr.; KF921668.1, JX870770.1, AF523073.1, AF522965.1, –; (S22) Ecuador, Ferm 50 (Q); MW849564*, –, –, MW940468*, –; *Samanea saman;* KX852445.1; *Samanea tubulosa* (Benth.) Barneby & J. W. Grimes; EF638135.1, EF638212.1, KX581235.1, –, –; ***Sanjappa** cynometroides* (Bedd.) E. R. Souza & Krishnaraj, KR997871.1, KR997866.1, –, KR997878.1, –; ***Senegalia*** Raf. sp. (S21) Ecuador, Ferm 31 (Q); MW849569*, MZ015543*, MZ169634*, MW940470*, *ycf1*\*; *Senegalia senegal* Britton; EF638152, HQ605075, KY688934, AF522976, –; *Senegalia westiana* Britton & Rose (S23) Puerto Rico, Steve Maldonado Silvestrini 278 (UPRRP); –*, MZ015544*, –*, MW940471*, *ycf1*\*; ***Serianthes** nelsonii* Merr. –, –, KX302353.1, –, –; ***Thailentadopsis** nitida* (Vahl) G. P. Lewis & Schrire; KF921670.1, JX870772.1, KX581237.1, JX870888.1; ***Vachellia** farnesiana* (L.) Wight & Arn.; EF638128.1, AF360728.1, AY574103.1, AY574119.1, –; (S19) Ecuador, Ferm 35 (S); MW849551*, MZ015545*, –*, MW940472*, *ycf1*\*; ***Viguieranthus** ambongensis* (R. Vig.) Villiers; KR997873.1, JX870773.1, KX581238.1, JX870890.1, –; (V2) Madagascar, D. J. Du Puy et al. M726 (P) MW849540*, MZ015546*, –*, MW940473*, *ycf1*\*; *Viguieranthus* Villiers cf. *ambongensis* (S35) Madagascar, Martial et al. 233 (TAN) –, –, –, –, *ycf1*\*; *Viguieranthus brevipennatus* Villiers (V3) Madagascar, R. W. Bussmann et al. 15178 (P) MW849541*, MZ015547*, MZ169645*, MW940474*, *ycf1*\*; *Viguieranthus cylindriostachys* Villiers (V4) Madagascar, N. Dumetz 1397 (P) MW849542*, MZ015548*, MZ169646*, MW940475*, *ycf1*\*; *Viguieranthus densinervus* Villiers; KR997874.1, JX870774.1, –, JX870891.1, –; (S33) Madagascar, Randriauaivo et al 1763 (TAN) –, MZ015549*, –, MW940476*, *ycf1*\*; *Viguieranthus densinervus* var. *pubescens* Villiers, (V6) Madagascar, Rakotoson 10393 RN (P) MW849543*, MZ015550*, MZ169676*, MW940477*, *ycf1*\*; *Viguieranthus glaber* Villiers; –, JX870775.1, KX302357.1, JX870892.1, –, (V7) Madagascar, Service Forestier Madagascar 12049 SF (P) MW849544*, –, –, MW940478*, *ycf1*\*; *Viguieranthus glandulosus* Villiers (V8) Madagascar, R. Capuron 9096 bis–SF (P) MW849545*, MZ015551*, –, –, *ycf1*\*; *Viguieranthus kony* (R. Vig.) Villiers;, –, JX870777.1, –, –, –, (V9) Madagascar, Service Forestier Madagascar 13290 SF (P) MW849546*, –, –, –, –; *Viguieranthus longiracemosus* Villiers (V10) Madagascar, R. Capuron 27996 SF (P) –, –, –, MW940479*, *ycf1*\*; *Viguieranthus megalophyllus* (R. Vig.) Villiers; KR997875.1, JX870776.1, –, –, –; (V11) Madagascar, R. Rabevohitra 2354 (P) MW849536*, MZ015552*, MZ169640*, MW940480*, *ycf1*\*; *Viguieranthus perrieri* (R. Vig) Villiers (V12) Madagascar, M. Y. Ammann MYA 395 (P) MW849537*, MZ015553*, MZ169629*, MW940481*, *ycf1*\*; *Viguieranthus pervillei* (Drake) Villiers (V14) Madagascar, P. Ranirison 344 (P) MW849538*, MZ015554*, –, MW940482*, *ycf1*\*; *Viguieranthus subauriculatus* Villiers; KR997876.1, JX870778.1, –, –, –; *Viguieranthus umbilicus* Villiers (V18) Madagascar, R. Capuron 27740 bis–SF (TAN) MW849533*, MZ015556*, MW940483*, MZ169677*, *ycf1*\*; *Viguieranthus unifoliolatus* Villiers (V19) Madagascar, Service Forestier Madagascar 4990 SF (P) MW849534*, MZ015557*, MW940484*, MZ169637*, *ycf1*\*; *Viguieranthus variabilis* Villiers (V20) Madagascar, Jardin Botanique Tananarive 3320 (TAN) MW849535*, MZ015558*, –, MW940485*, –; *Viguieranthus* Villiers sp. (Mada125) –, –, KC479271.1, –, –; (S32) Madagascar, Du Puy et al M175 (TAN) MW849539*, MZ015555*, –, –, *ycf1*\*; ***Wallaceodendron** celebicum* Koord.; EF638097.1, EF638222.1, KX570946.1, –, –; ***Zapoteca** aculeata* (Spruce ex Benth.) H. M. Hern. (K1) Ecuador, Delinks 332 (NY); MK622329, MK638924, –, MK622373, *ycf1*\*; (K2) MK622330, –, –, MK622363, –; *Zapoteca amazonica* (Benth.) H. M. Hern. (K30) Peru, Mexia 8295 (S); MK622344, MK638946, MZ169639*, MK622377, *ycf1*\*; *Zapoteca alinae* H. M. Hern.; –, JX870779.1, JX870893.1; (K3) Mexico, Pascual 1492 (NY); MK622336, MK638925, MZ169643*, MK622368, *ycf1*\*; (K4) Mexico, Gomez 91–7–7 (NY); MT926005, MK638926, MZ169647*, MT926027, *ycf1*\*; *Zapoteca andina* H. M. Hern.; –, MT937169, –, MT926024, –; (S38); –, –, –, MT926025, –; (S39) Ecuador, Ståhl L102 (S); –, MT937170, MZ169644*, MT926026, –; *Zapoteca balsasensis* H. M. Hern., Mexico, Contreras & Thomas 1735 (NY); MT926006, MK638928, MZ169648*, MT926023, *ycf1*\*; *Zapoteca caracasana* (Jacq.) H. M. Hern. subsp. *caracasana* (K34) Hispaniola, Ekman 16527 (S); MK622335, MK638949, MZ169649*, MK622370, *ycf1*\*; *Zapoteca caracasasa* subsp. *weberbaueri* (Harms.) H. M. Hern. (K32) Ecuador, Asplund 15982 (S); MK622345, MK638947, –, MK622374, *ycf1*\*; (K33) Ecuador, Asplund 15503 (S); MK622333, MK638948, MZ169650*, MK622376, *ycf1*\*; (S40) Ecuador, Ståhl L101 (S); MT926015, MT937168, MZ169651*, –,*ycf1*\*; *Zapoteca costaricensis* (Britton & Rose) H. M. Hern. (513) –, MK638961, –, –, –; *Zapoteca cruzii* H. M. Hern. (505) Mexico, Gual 272 (MEXU); MK622328, MK638962, MZ169652*, MK622375, *ycf1*\*; *Zapoteca filipes* (Benth.) H. M. Hern.; –, JX870780.1, –, JX870896.1, –; (K9) –, MK638927, –, MK622367, –; *Zapoteca formosa* (Kunth.) H. M. Hern.; –, JX870781.1, –, JX870897.1, –; (K36) Novara & Bruno 8865 (S); –, MK638950, MZ169653*, MK622356,*ycf1*\*; *Zapoteca formosa* subsp. *formosa* (Kunth.) H. M. Hern. (K10) Mexico, McVaugh 20327 (NY); –, MK638929, MZ169654*, MT926022, *ycf1*\*; (K11) Mexico, McVaugh 19857 (NY); –, MK638930, MZ169655*, –, –; *Zapoteca formosa* subsp. *mollicula* (J.F.Macbr.) H. M. Hern. (K13) Mexico, Hughes 1804 (NY); MK622337, MK638931, MZ169658*, MK622362, *ycf1*\*; *Zapoteca formosa* subsp. *rosei* (Wiggins) H. M. Hern. (K15) MK622353, MK638932, –, –, –; *Zapoteca formosa* subsp. *salvadorensis* (Britton & Rose) H. M. Hern. (K16) MK622352, MK638933, –, –, –; (K17) Guatemala, Williams & al. 22456 (NY); MK622338, MK638934, MZ169659*, MK622355, *ycf1*\*; *Zapoteca formosa* subsp. *schottii* (Torr. ex S. Watson) H. M. Hern. (K18) US/Arizona, Parker 5861 (NY); MT926014, MK638935, MZ169661*, –, *ycf1*\*; (K19) US/Arizona, Kearney & Peebles 14960 (NY); MK622339, MK638936, MZ169662*, MK622357, *ycf1*\*; (532) Semillas cultivadas XDL89–0405D (CICY); MK638923, MT937167, MZ169660*, MK622379, *ycf1*\*; *Zapoteca formosa* subsp. *socorrensis* (I. M. Johnst.) G. A. Levin, H. M. Hern. & Moran (K20) Mexico, Moran 25546 (NY); MK622340, MK638937, –, –, *ycf1*\*; *Zapoteca gracilis* (Griseb.) Bässler (K37) Bahamas, Howard 10021 (S); MK622346, MK638951, MZ169656*; MK622362, *ycf1*\*; (K38) Cuba, Ekman 8198 (S); MK622347, MK638952, MZ169657*, MK622359, *ycf1*\*; (K39) Bahamas, Webster, Samule & Williams 10511A (S); MK622348, MK638953, MZ169627*, MK622372, –; *Zapoteca lambertiana* (G. Don) H. M. Hern.; –, JX870782.1, –, JX870894.1, –; (K21) Mexico, Breedlove 36732 (NY); MK622331, MK638938, MZ169663*, MK622360, *ycf1*\*; (K22) MK622332, MK638939, –, –, –; *Zapoteca media* (M. Martens & Galeotti) H. M. Hern. (K23) Mexico, Moore Jr. 3986 (NY); MK622341, MK638940, MZ169630*, MK622365, *ycf1*\*; (K24) Mexico, Johnston 12043 (NY); MK622351, MK638941, MZ169664*, MK622366, *ycf1*\*; *Zapoteca microcephala* (Britton & Killip) H. M. Hern. Colombia, Haught 1711 (K); MT926013, MT937166, MZ169665*, MT926021, ycf1*; *Zapoteca mollis* (Standl.) H. M. Hern.; –, –, JQ587906, –, –; (K25) Costa Rica, Rodriguez 2420 (NY); MK622342, MK638942, MZ169666*, MT926019, –; (K26) Costa Rica, Grayum 4201 (NY); MT926012, MT937165, –, MT926020, *ycf1*\*; *Zapoteca nervosa* (Urb.) H. M. Hern. (K40) MK622349, MK638954, –, –, –; (K41) Hispaniola, Ekman 15423 (S); MT926011, MK638955, –, MK622358, *ycf1*\*; *Zapoteca portoricensis* subsp. *flavida* (K) MT926010, –, –, –, –; *Zapoteca portoricensis* (Jacq.) H. M. Hern.; –, –, KJ012829, –, –; *Zapoteca portoricensis* subsp. *portoricensis* (K42) Hispaniola, Ekman 10924 (S); MK622350, MK638956, MZ169631*, MK622371, *ycf1*\*; (K43) Hispaniola, Ekman 13376 (S); MK622334, MK638957, MZ169622*, MT926017, *ycf1*\*; (S28) Puerto Rico, Ferm 105 (UPRRP); MT926009*, MT937164*, –, MT926018*, –; *Zapoteca portoricensis* subsp. *pubicarpa* H. M. Hern. (K27) Mexico, Purpus 2668 (NY); MT926008, MK638943, MZ169667*, MT926016, –; *Zapoteca quichoi* H. M. Hern. & A. M. Hanan (498) Mexico, Calónico 21109 (MEXU); MK622327, MK638960, MZ169621*, MK622364, *ycf1*\*; *Zapoteca ravenii* H. M. Hern. (K28) Mexico, Martinez 23967 (NY); MK622343, MK638944, MZ169668*, MK622369, *ycf1*\*; *Zapoteca scutellifera* (Benth.) H. M. Hern. (K29) Brazil, Amaral 1231 (NY); –, MK638945, –, –, *ycf1*\*; *Zapoteca sousae* H. M. Hern. & A. Campos; –, JX870783.1, KX581240.1, –, –; *Zapoteca tehuana* H. M. Hern. (339) Mexico, A. Campos 4108 (MEXU); MK622326, MK638963, MZ169620*, MK622378, *ycf1*\*; (338) Mexico, Torres Colín 8934 (MEXU); MT926007, MK638959, MZ169669*, MK622354, *ycf1*\*; *Zapoteca tetragona* (Willd.) H. M. Hern. (K45) –, MK638958, –, –, –; ***Zygia** basijuga* (Ducke) Barneby & J. W. Grimes (S10) Ecuador, Ferm 60 (Q); MW849524*, MZ015537*, MZ169625*, MW940463*, *ycf1*\*; (S11) Ecuador, Ferm 71 (Q); MW849525*, MZ015538*, MZ169675*, MW940464*, *ycf1*\*; (S12) Ecuador, Ferm 88 (Q); MW849526*, MZ015539*, MZ169626*, MW940465*, *ycf1*\*; *Zygia latifolia* (L.) Fawc. & Rendle (S26) Jamaica, Ferm 103 (IJ); MW849527*, –, –, –, –; (S27) Jamaica, Ferm 104 (IJ); MW849528*, MZ015559*, MZ169670*, MW940486*, *ycf1*\*; *Zygia racemosa* (Ducke) Barneby & J. W. Grimes; –, –, JQ626423.1, –, –; (J16); MK681163, MK641678, –, MK903295, –; *Zygia* P. Browne sp. (S6) Ecuador, Ferm 37 (Q); MW849531*, MZ015561*, MZ169672*, MW940488*, *ycf1*\*; (S7) Ecuador, Ferm 15 (Q); MW849532*, MZ015562*, MZ169673*, MW940489*, *ycf1*\*; (S8) Ecuador, Ferm 55 (Q); MW849523*, MZ015563*, –, MW940490*, –; (S30) Ecuador, Ferm 23 (Q); MW849529*, MZ015560*, MZ169671*, –, –; (S31) Ecuador, Ferm 24 (Q); MW849530*, –, –, MW940487*, –;

