## Supplementary figures and images for "Phylogeny of the ingoid clade (Caesalpinioideae, Fabaceae), based on nuclear and plastid data"

### supplementary figure 1

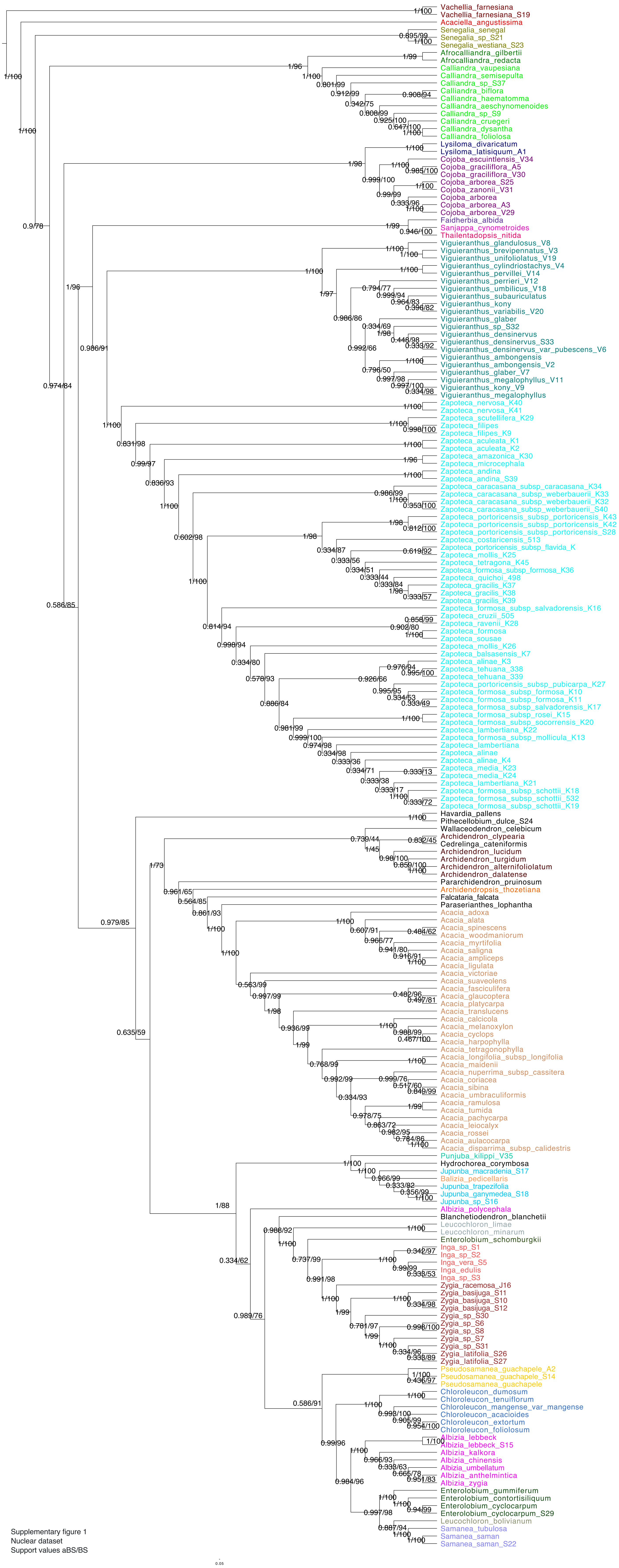


Supplementary figure 1
Nuclear dataset
Support values aBS/BS

### supplementary figure 2

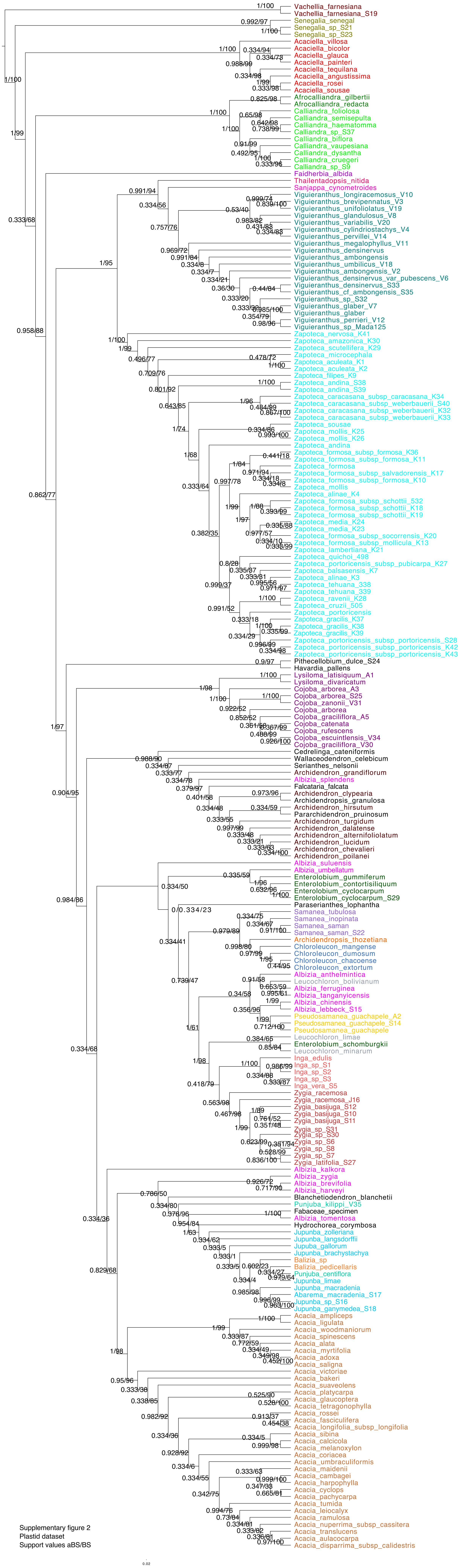


Supplementary figure 2
Plastid dataset
Support values aBS/BS
