## supplementary figure 3 for "Phylogeny of the ingoid clade (Caesalpinioideae, Fabaceae), based on nuclear and plastid data"

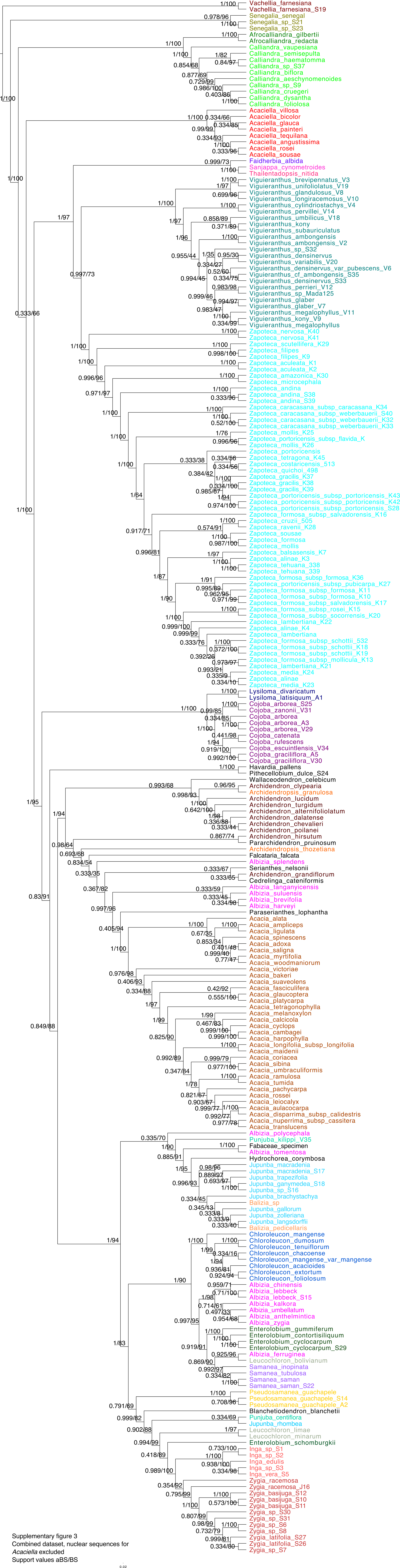


Supplementary figure 3
Combined dataset, nuclear sequences for
*Acaciella* excluded
Support values aBS/BS
